# Regenerative cell turnover coupled to a tumor-hostile niche protects naked mole-rats from cancer

**DOI:** 10.1101/2023.10.10.561212

**Authors:** Mikaela Behm, Pablo Baeza-Centurión, Sabrina Weser, Veronica Francis Busa, Pietro Zafferani, Luca Penso-Dolfin, Javier Botey-Bataller, Sylvain Delaunay, Nick Hirschmüller, Beatrice Berthet, Marie-Luise Koch, Stefania Del Prete, Milica Bekavac, Daniela Sohn, Liana Penteskoufi, Celine Reifenberg, Francesca Coraggio, Meike Schopp, Fritjof Lammers, Llorenç Solé-Boldo, Jeleana Dutton, Rebecca E. Wagner, Sandra Blanco, Anke S. Lonsdorf, Sabine Begall, Walid T. Khaled, Ewan St. J. Smith, Duncan T. Odom, Angela Goncalves, Michaela Frye

## Abstract

Long-lived species suppress cancer despite accumulating somatic mutations throughout life, but how this is achieved in renewing tissues remains unclear. Here, we show that naked mole-rat skin uncouples high cellular turnover from cancer risk through coordinated epithelial and stromal mechanisms that constrain clonal outgrowth and suppress tumor-promoting inflammation. Despite elevated epidermal proliferation and mutational burden, naked mole-rat skin maintains tissue integrity through an expanded pool of early committed progenitor (hybrid) cells and replication-coupled genome maintenance pathways. Under chronic carcinogenic stress, undifferentiated basal cells replenish the hybrid progenitor pool. This dilutes initiated clones and permits only limited selective expansion of rare cancer gene–mutant clones. Depletion of the hybrid compartment shifts this protective state toward clonal expansion and inflammatory activation. These expanding clones are further constrained by a highly tumor-suppressive stromal microenvironment, driven by fibroblasts that adopt a metabolically restricted, non-inflammatory program. Together, our data uncover a multi-layered tumor-suppressive strategy that couples turnover-driven regeneration with an anti-permissive stromal niche to prevent malignant progression under mutagenic stress.

## Introduction

Across mammals, cancer incidence is not explained by somatic mutation burden or proliferation alone. Long-lived species must sustain tissue integrity over decades of cumulative genotoxic exposure and continuous cellular turnover. To avoid cellular transformation, tissue-intrinsic tumor-suppressive programs buffer oncogenic variation and constrain malignant progression. Consistent with this, comparative sequencing across mammals shows that somatic mutation rates per year scale inversely with lifespan, such that end-of-life mutation burdens are far more similar across species than their lifespans and body sizes would predict ^1^. For example, the mouse-sized naked mole-rat (*Heterocephalus glaber*) with exceptional longevity (>35 years) and high natural cancer resistance exhibits an annual somatic mutation rate similar to that of the giraffe, which also has a lifespan of ∼35-years ^1–4^.

Malignant transformation is exceptionally rare in naked mole-rats. Among thousands of captive animals fewer than ten neoplasms have been observed ^5–8^, although naked mole-rat cells can be transformed in the presence of oncogenes both *in vitro* and *in vivo* ^9–11^. The mechanisms underlying their exceptional cancer resistance remain poorly understood. Both cell autonomous and non-autonomous mechanisms have been reported, including enhanced DNA repair ^12–15^, heightened sensitivity to contact-inhibition ^16–19^, distinct activation of tumor-suppressors ^20^, and a lack of or dampened immune cell response ^21–23^. How these processes are integrated with stem-progenitor dynamics, tissue architecture and cell fate regulation remains largely unclear.

Disruptions in tightly balanced stem cell self-renewal and differentiation processes must be prevented, eliminated, or functionally compensated to preserve healthy tissue regeneration throughout life ^24^. Harmful mutations that either promote excessive cellular expansion or impair regenerative capacity can ultimately lead to cancer or premature aging, respectively ^25^. Despite growing attention to naked mole-rat cancer resistance, it remains unresolved whether protection reflects uniformly reduced mutational input across tissues, or whether certain organs tolerate substantial mutagenesis without increased cancer risk. As a rapidly renewing barrier that is persistently challenged by genotoxic injury, the epidermis provides a tractable system to interrogate how tissues tolerate or eliminate mutational damage. Here, we directly test how naked mole-rat skin maintains tissue integrity under chronic carcinogenic exposure, and more broadly, how cell state and tissue architecture buffer mutational burden to constrain malignant progression.

By integrating mutational profiling with single-cell and perturbation-based analyses, we uncover that epidermal lineage dynamics together with the dermal microenvironment jointly constrain clonal expansion under mutagenic stress. Despite substantial mutational burden and high cellular turnover, chronic carcinogenic exposure fails to disrupt tissue integrity in naked mole-rats. Rather than relying on widespread cell loss, skin homeostasis is maintained by cell state-coupled genome maintenance pathways, turnover-driven clonal dilution and a non-permissive stromal niche that collectively constrain clonal evolution.

## Results

### Turnover-driven clonal expansion is uncoupled from cancer risk in naked mole-rat skin

Across mammals, naked mole-rats show the fewest mutations per genome in clonal intestinal crypts ^1^. It remains unknown whether a similar relationship holds in skin, where tissue integrity is sustained by multiple distinct tissue-specific stem cell populations, each driving its own turn-over ^26^. Therefore, we performed high-depth whole-exome sequencing on a spatially resolved grid of 1 mm biopsies from naked mole-rats and mice (Fig. 1a; Extended Data Fig. 1a). We observed slightly higher somatic mutation rates in naked mole-rat skin biopsies than in mouse, along with a significant increase in potentially harmful mutations, frequently affecting genes linked to malignant transformation (Fig. 1b-c; Extended Data Fig. 1b-d). The elevated mutational burden correlated with larger clonal expansions across the 4 × 4 mm skin area, as reflected by a higher fraction of variants with allele frequencies (VAFs) greater than 0.1 in naked mole-rats (Fig. 1c-f; Extended Data Fig. 1d-f).

**Fig. 1:**
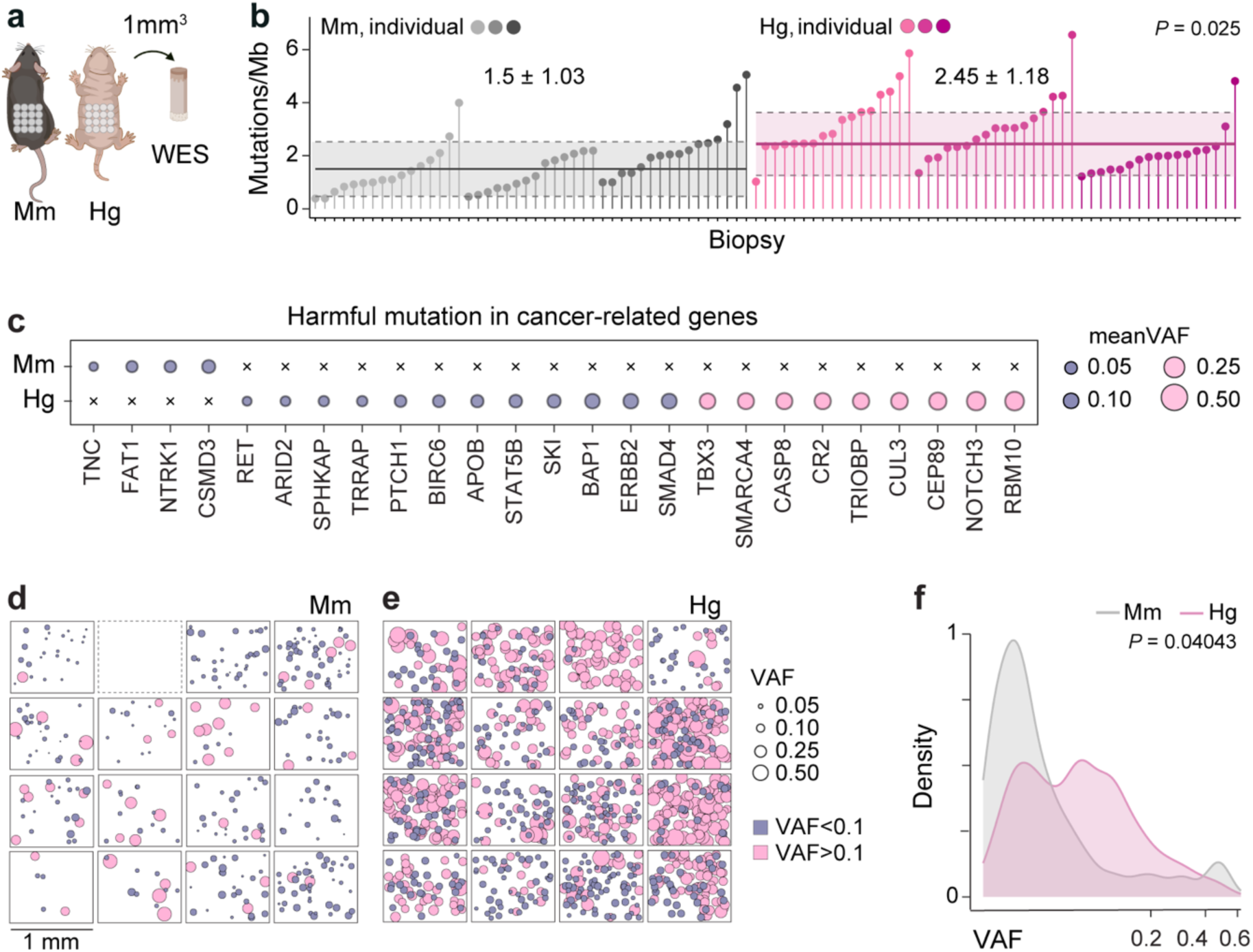
Somatic mutational landscapes in cancer-permissive and- resistant skin. **a**, 1 mm³ full-thickness dorsal skin biopsies from mouse (*Mm*) and naked mole-rat (*Hg*) collected in a 4 x 4 grid for whole-exome sequencing. **b**, SNV burden in *Mm* and *Hg* skin biopsies grouped by individual and shaded by species. Solid lines indicate the median; dotted lines indicate ± s.d. **c**, Harmful mutations in cancer-associated genes in *Mm* and *Hg* skin. Filled circles indicate mutated genes, with size proportional to mean VAF; crossed symbols indicate absence of mutations. **d–e**, Nonsynonymous SNVs with predicted moderate or high functional impact in *Mm* (**d**) and *Hg* (**e**) biopsies. Each square represents one biopsy; each circle represents one mutated gene; circle size indicates VAF (blue, <0.1; pink, >0.1). Dotted square indicates no mutated gene passing filtering criteria. **f**, Distribution of harmful nonsynonymous SNVs in *Mm* and *Hg* skin shown on a pseudo-logarithmic scale. N, individuals; n, biopsies. *Mm*, N = 3, n = 14–16; *Hg*, N = 3, n = 17 (**b**, **c**, **f**). *Mm*, N = 1, n = 16; *Hg*, N = 1, *n* = 16 (**d**, **e**). Linear mixed-effects model (**b**). One-sided Wilcoxon rank-sum test (**f**).

Larger mutant clones could reflect either increased tissue turnover, which accelerates clonal expansion, or reduced turnover, which prolongs clonal persistence. To directly assess cellular turnover, we measured the proliferative capacity of the interfollicular epidermis (IFE), which is maintained by stem cells located in the basal layer of the epithelium ^27^. To our surprise, we found that naked mole-rat skin has a two-fold higher proliferation rate in the IFE compared to both human and mouse (Fig. 2a-b). A combination of fast-cycling progenitors and oncogenic mutations increases cancer risk ^28^, except when compensatory cellular mechanisms are in place that either limit self-renewal and cycling or promote differentiation.

**Fig. 2:**
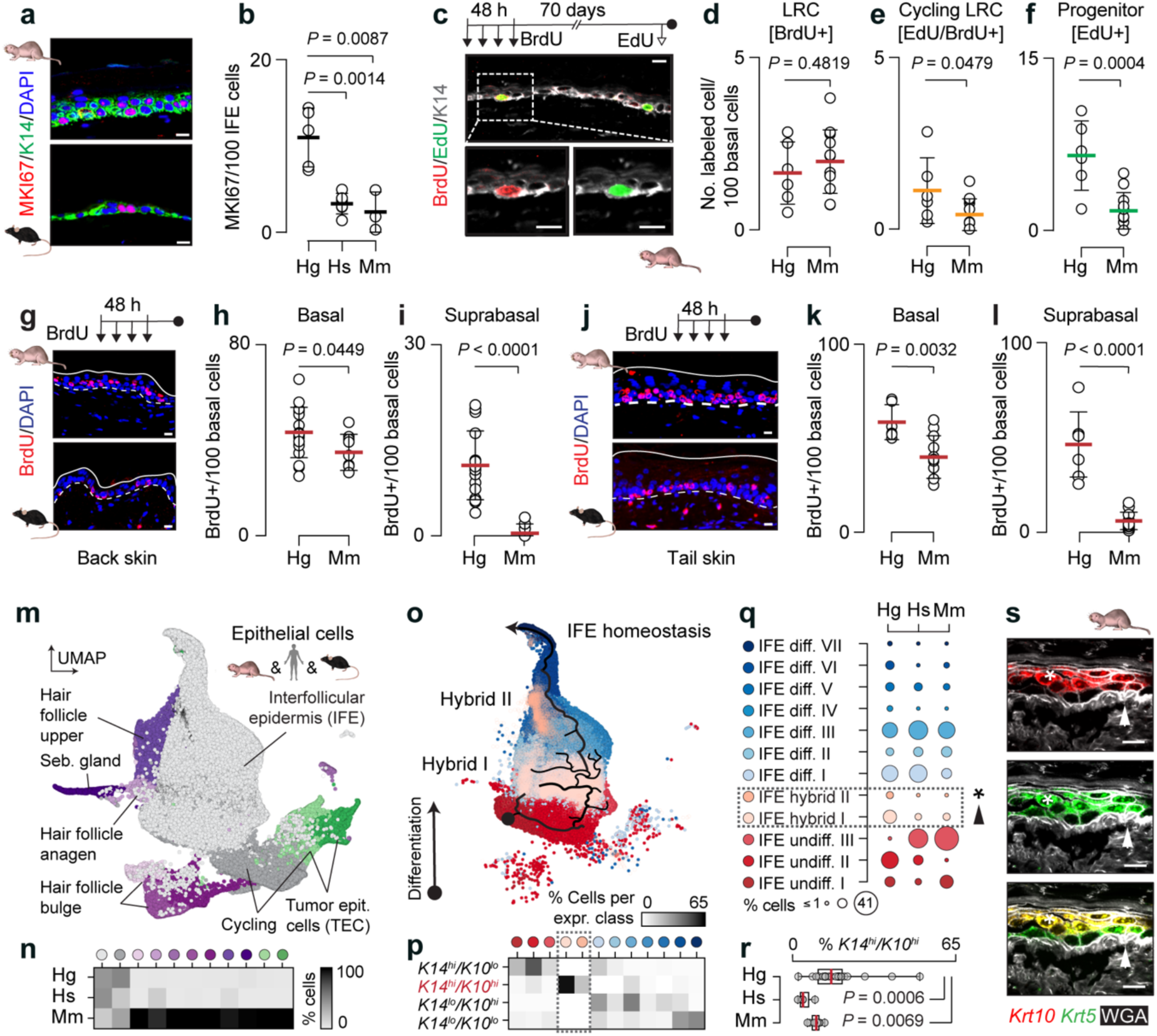
Rapid epidermal homeostatic regeneration in naked mole-rat skin. **a**, **b**, Immunofluorescence (**a**) and quantification (**b**) of MKI67*^+^*cells in homeostatic naked mole-rat (*Hg*), human (*Hs*), and mouse (*Mm*) IFE. Data points represent biological replicates (N = 3–5). **c–f**, Immunofluorescence (**c**) and quantification (**d–f**) of BrdU^+^ (**c**, **d**), EdU^+^/BrdU^+^ (**c**, **e**), and EdU^+^ (**f**) cells in the basal layer of *Hg* and *Mm* back skin. LRC, label-retaining cell. **g–l**, Immunofluorescence (**g**, **j**) and quantification of basal (**h**, **k**) and suprabasal (**i**, **l**) BrdU^+^ cells in *Hg* and *Mm* back (**g–i**) and tail (**j–l**) skin. Data points represent individual measurements from three naked mole-rats and three mice (**d–f**, **h**, **i**, **k**, **l**). **m**, UMAP of *Hg*, *Hs*, and *Mm* epithelial cells. **n**, Weighted average % epithelial cell types per species. **o**, UMAP of *Hg*, *Hs*, and *Mm* IFE cells with an overlaid pseudotime differentiation trajectory, starting at the black dot. Colours indicate cell differentiation states. **p**, % IFE cells with high or low *K14* and *K10* expression. **q**, % of IFE cell state per species. **r**, % of K14^hi^/K10^hi^ cells per skin biopsy, by species. *Hg*, N = 6, n = 21; *Hs*, N = 5, n = 5; *Mm*, N = 3, n = 9 (**m–r**). **s**, RNAscope showing *Krt10* and *Krt5* expression in *Hg* skin. WGA, plasma membrane label. Arrowhead, IFE hybrid I; asterisk, IFE hybrid II. Scale bars, 10 μm (**a**, **c**, **g**, **j**, **s**). Box plot shows median and interquartile range, with whiskers extending to minimum and maximum values (**r**). Values show mean ± s.d. (**b**, **d–f**, **h**, **k**, **l**) and median with interquartile range (**i**). Unpaired two-tailed *t*-test (**b**, **d–f**, **h**, **i**, **k**, **l**, **r**). Two-tailed Mann–Whitney U test (**i**). Exact *P* values are shown.

### Precocious differentiation ensures tissue integrity despite high cellular turnover

To measure how stem cell quiescence is balanced with progenitor proliferation, we performed label-retaining cell (LRC) assays, which identify long-lived, slow-cycling cells (BrdU^+^) over time (Fig. 2c) ^29^. Actively cycling cells were concurrently labelled by administering a single EdU pulse shortly before sample collection (Fig. 2c). Neither treatment altered skin morphology (Extended Data Fig. 2a-b). While we observed a similar number of quiescent cells (Fig. 2d), naked mole-rat epidermis contained a larger fraction of actively cycling LRCs and EdU^+^ progenitors (Fig. 2e-f). Similarly, a 48-hour chase revealed significantly more BrdU⁺ cells in the basal and suprabasal layers of naked mole-rat epidermis (Fig. 2g-i). Since the number of epidermal layers differs in back skin between naked mole-rat and mouse (Extended Data Fig. 2a-b), we independently validated the higher proliferation rates in tail skin, which displays a comparable multilayered epithelium in both species (Fig. 2j-l). In conclusion, fast progenitor cycling in naked mole-rats was not counterbalanced by fewer quiescent or slower cycling cells. Instead, naked mole-rat epidermis was highly regenerative in homeostasis, which was further supported by elevated expression of keratin 6 (KRT6), a marker of wound-induced regeneration in mouse and human (Extended Data Fig. 2c-f) ^30^.

We next investigated whether changes in differentiation dynamics might maintain tissue integrity. We performed single-cell RNA sequencing and determined epidermal progenitor fates in the IFE. First, we integrated and clustered single-cell transcriptomes from naked mole-rat, human and mouse, and annotated all major epidermal compartments (Fig. 2m; Extended Data Fig. 2g). Then, we removed sebaceous-gland and hair-follicle populations (Fig. 2n; coloured cells). Finally, we performed trajectory analysis revealing a continuous progression from undifferentiated K14⁺ to committed K10⁺ progenitors and terminally differentiated cells (Fig. 2o; black line). This progression was interrupted only by short branches of hybrid cells co-expressing K14/K10 (Hybrid I), mostly consisting of naked mole-rat epidermal cells (Fig. 2o-r, light beige colour). We observed a second hybrid state (Hybrid II), which was more differentiated and positioned further along the trajectory (Fig. 2o-q, dark beige). We confirmed in naked mole-rat and mouse skin that keratin markers for undifferentiated (*Krt14* or *Krt5*) and differentiated (*Krt10*) keratinocytes were indeed co-expressed within the same cells in both the basal and suprabasal epidermal layers (Fig. 2s; Extended Data Fig. 2h-i).

This enriched hybrid population in the basal layer of the epidermis represents cells committed to differentiation, with proliferation strictly limited to only a few rounds of division ^31–33^. Epidermal cells simultaneously expressing progenitor- and differentiation-related keratins have been reported in newborn and adult mice, and neonatal human foreskin ^31,34,35^; however, we found that their localisation in the basal layer was extremely rare when compared to naked mole-rat skin (Extended Data Fig. 2j-k). Thus, naked mole-rats maintain epidermal homeostasis by balancing fast-cycling progenitors with an enlarged pool of committed cells, thereby sustaining differentiation while preventing tissue overgrowth.

### A larger population of committed progenitors efficiently buffers damage to basal cells

To assess the functional contribution of basal-layer committed progenitors to the maintenance of epidermal integrity, we exposed the skin to the mutagen DMBA for 48 hours (Fig. 3a; top). Following acute damage, the size of the hybrid population significantly decreased in both species, consistent with the hypothesis that it serves as a buffer, being preferentially lost through terminal differentiation in response to injury (Fig. 3a-b; Extended Data Fig. 3a-b). In addition, mice showed premature differentiation of basal progenitors with a significant increase in suprabasal K14^+^/K10^+^ cells, whereas naked mole-rat suprabasal K14^+^/K10^+^ cell numbers remained unchanged (Fig. 3a, c). We reasoned that naked mole-rat basal epidermal cells either completed differentiation without ‘stalling’ in a hybrid state or were efficiently eliminated through cell death. Therefore, we next quantified damaged and apoptotic cells.

**Fig. 3:**
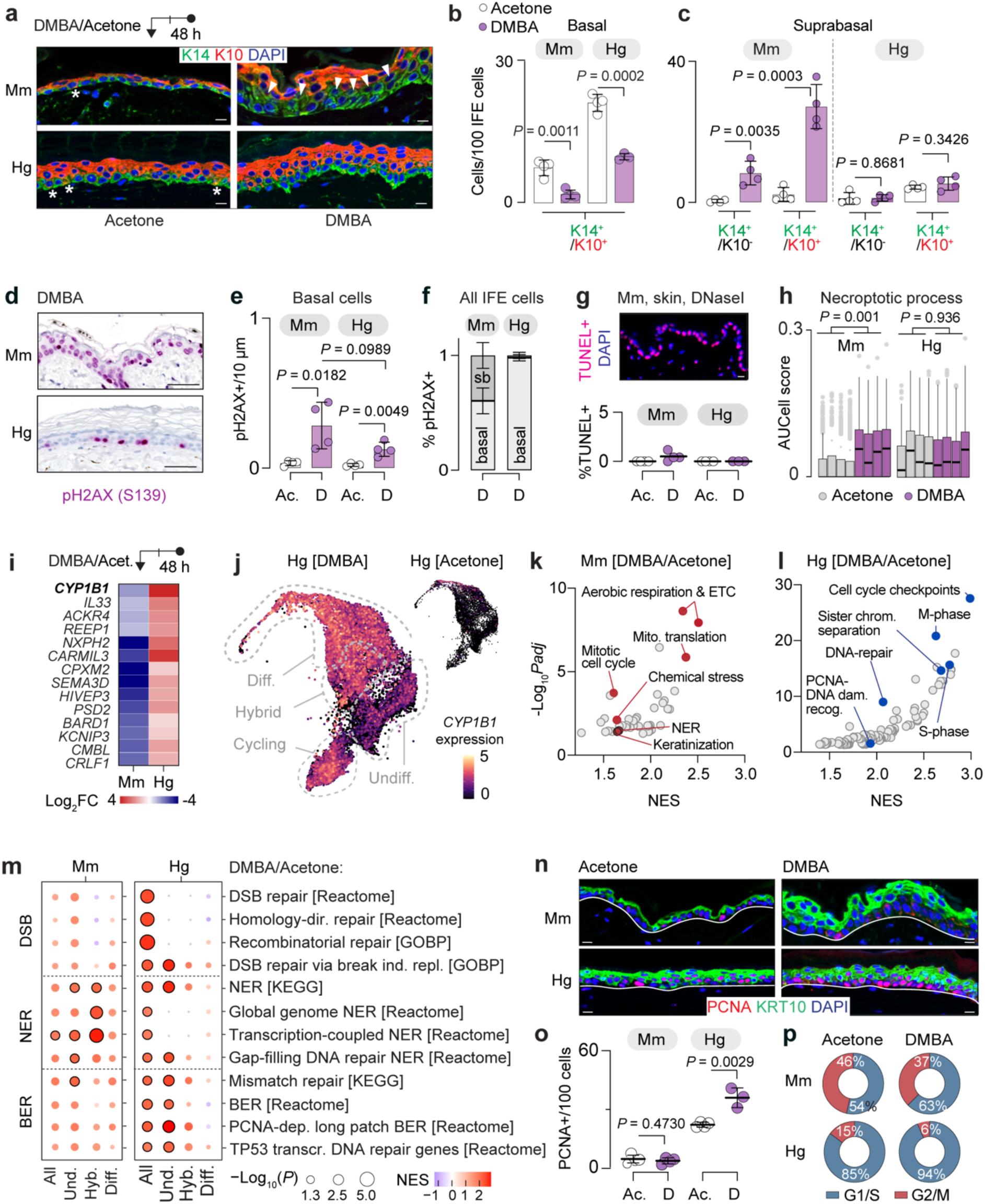
Mouse resolves DNA damage by delamination, naked mole-rat by genome maintenance. **a–c**, Immunofluorescence (**a**) and quantification of basal (**b**) and suprabasal (**c**) K14^+^ and K10^+^ IFE cells in mouse (*Mm*) and naked mole-rat (*Hg*) skin 48 h after a single topical application of DMBA or acetone. Asterisks mark K10⁺ basal cells; arrowheads mark suprabasal K14⁺/K10⁻ cells (**a**). **d–f**, Immunohistochemistry (**d**) and quantification of pH2AX (S139)⁺ cells per 10 μm basal footprint (**e**) and distribution of pH2AX (S139)^+^ cells in basal and suprabasal (sb) layers (**f**) in *Mm* and *Hg* acetone (Ac.)- and DMBA (D)-treated skin. **g**,TUNEL immunofluorescence (top) and quantification per 100 IFE cells (bottom) in Ac.- or D-treated *Mm* and *Hg* skin; DNase I–treated *Mm* skin serves as positive control. **h**, Gene set scoring of the GOBP *necroptotic process* in undifferentiated IFE cells. **i**, Log₂ fold change of significant (*P*adj < 0.05) differentially regulated genes in *Mm* and *Hg* IFE cells 48 h after DMBA treatment relative to acetone control. **j**, CYP1B1 expression projected onto a UMAP of *Hg* IFE cells treated with DMBA or acetone. **k**, **l**, Gene set enrichment analysis showing significant enriched pathways (*P*adj < 0.05) in *Mm* and *Hg* IFE cells comparing DMBA with acetone. **m**, GSEA of DNA repair pathways in *Mm* and *Hg* epithelial cell states; dot size indicates −log₁₀(*P*adj), colour indicates NES, black outlines mark significant (*P*adj < 0.05) comparisons. **n**, **o**, Immunofluorescence (**n**) and quantification (**o**) of PCNA⁺ basal IFE cells in *Mm* and *Hg* skin 48 h after Ac. or D treatment. **p**, Cell-cycle distribution of undifferentiated IFE cells. Scale bars, 10 μm (**a**, **g**, **n**), 50 μm (**d**). Values show mean ± s.d. (**b**, **c**, **e–g**, **o**). Data points represent individual means (**b**, **c**, **e–g**, **o**). Box plots show median and interquartile range, with 1.5× IQR whiskers; each box represents one individual (**h**). N = 3–4 biologically independent individuals per species and treatment (**b**, **g**, **o**); N = 4 biologically independent individuals per species and treatment (**c**, **e**, **f**, **h–m**). Unpaired two-tailed *t*-test (**b**, **c**, **e**, **o**). Linear mixed-effects model (**h**). Exact *P* values are shown.

To visualize damaged cells in the epidermis, we labelled DNA double-strand breaks by detecting phosphorylated γH2AX (Fig. 3d; Extended Data Fig. 3c-e). While both species accumulated γH2AX foci in the basal layer, only mice additionally accumulated damage in suprabasal keratinocytes (Fig. 3d-f; Extended Data Fig. 3d-e). Thus, mice cleared damaged progenitors through delamination and terminal differentiation (Extended Data Fig. 4a-b). To test whether naked mole-rats instead eliminated damaged cells through apoptosis, we performed TUNEL assays to detect DNA fragmentation (Fig. 3g; Extended Data Fig. 4d-e). Apoptosis was not increased in either species, but mouse epidermis up-regulated necroptosis as expected (Fig. 3h) ^21,36^. Since the absence of TUNEL-positive cells did not exclude alternative mechanisms for expelling damaged cells, we performed single-cell transcriptome analyses to search for other cell death-related pathways. In contrast to mouse, naked mole-rat epidermis did not engage in any form of cell death (Extended Data Fig. 4c, f-j). We concluded that, in response to acute DNA damage, naked mole-rat epidermis eliminated cells neither through premature differentiation nor cell death.

### Genome maintenance, not differentiation, shields naked mole-rat skin from acute damage

Since naked mole-rat epidermis retained damaged cells under stress, we reasoned that protective mechanisms must be engaged to limit the accumulation of genomic lesions. To test whether DMBA exposure elicits species-specific protective responses, we first assessed DMBA mutagenicity *in vitro*. Whole-genome sequencing of DMBA-treated clonal dermal fibroblasts revealed comparable mutagenic activity in both species (Extended Data Fig. 4k-l). We then performed single-cell transcriptome analyses to identify repair pathways induced by acute DMBA exposure (Fig. 3i–l). Naked mole-rat epidermis responded to the genotoxic stress by strongly inducing CYP1B1, a key enzyme that converts DMBA into DNA-reactive metabolites ^37^, in addition to other genes associated with repair programs, including metabolic and detoxification pathways (CMBL) ^38^, DNA damage sensing and repair (BARD1) ^39^, and stress adaptation, immune regulation, and tissue remodelling (IL33, ACKR4, SEMA3D, CARMIL3, CPXM2, CRLF1) (Fig. 3i-j; Extended Data Fig. 5a-r). Indeed, only naked mole-rats specifically up-regulated DNA double-strand break-related repair pathways in response to DMBA-induced genotoxic stress (Fig. 3m; top panels).

Although both naked mole-rat and mouse activated repair pathways in response to DMBA, they relied on different protective mechanisms (Fig. 3k-m). In mice, DMBA induced keratinocyte differentiation-associated signatures together with mitochondrial activation (Fig. 3k) ^40^. As differentiated epidermal cells are metabolically and transcriptionally active but non-proliferative, the hybrid population (Hyb.) in mice showed a relative increase in transcripts associated with transcription-coupled nucleotide excision repair (NER), suggesting a shift in repair toward transcriptional integrity over genome maintenance (Fig. 3m; middle panels).

In naked mole-rats, DMBA primarily activated replication stress-associated maintenance pathways that included cell-cycle checkpoint activation and S-phase DNA-damage recognition (Fig. 3m). In undifferentiated cells, we observed upregulation of transcripts associated with mismatch repair and base excision repair (BER) (Fig. 3m, bottom panel). In agreement with this finding, only naked mole-rat cells upregulated PCNA, a key coordinator of DNA synthesis and repair (Fig. 3n-o). Moreover, in naked mole-rats, checkpoint activation was accompanied by a reduction in the proportion of cells in G2/M and a prolonged S-phase, suggesting that replication-coupled repair processes are active during this interval (Fig. 3p). Naked mole-rats also showed a modest increase in the proliferation marker KI67 (Extended Data Fig. 5s-t). However, the total number of basal-layer cells remained unchanged, suggesting that this reflected checkpoint-mediated cell-cycle activity and loss of the hybrid population rather than increased proliferation (Extended Data Fig. 5u). Together, our data showed that, unlike in mice, naked mole-rat epidermal cells preferentially engaged genome-maintenance DNA repair pathways following DMBA exposure.

In summary, only naked mole-rats efficiently activated DNA repair pathways across all epidermal cell layers following DMBA exposure. Damage repair in the absence of cell death further implied that immune cells remained largely quiescent or non-inflammatory in this context. Therefore, we next studied how immune cells responded to acute DNA damage.

### Restraining the immune compartment preserves tissue integrity in naked mole-rats

In mammalian skin, immune cells are primarily found in the dermis ^41^. Our histological analyses revealed that the overall dermal cellularity was substantially lower in naked mole-rats and remained unchanged after exposure to DMBA (Fig. 4a-b). While naked mole-rats lack natural killer cells ^22^, their skin also showed comparatively low numbers of myeloid (macrophages, neutrophils, Langerhans cells, and conventional dendritic cells) and lymphoid (T cells and innate lymphoid cells) cells identified by scRNA sequencing and immunofluorescence staining for CD45 (Fig. 4c-e; Extended Data Fig. 6a-b) ^42^. Only mouse skin showed a significant increase in myeloid and lymphoid CD45^+^ immune cells in response to DMBA treatment (Fig. 4d-e).

**Fig. 4:**
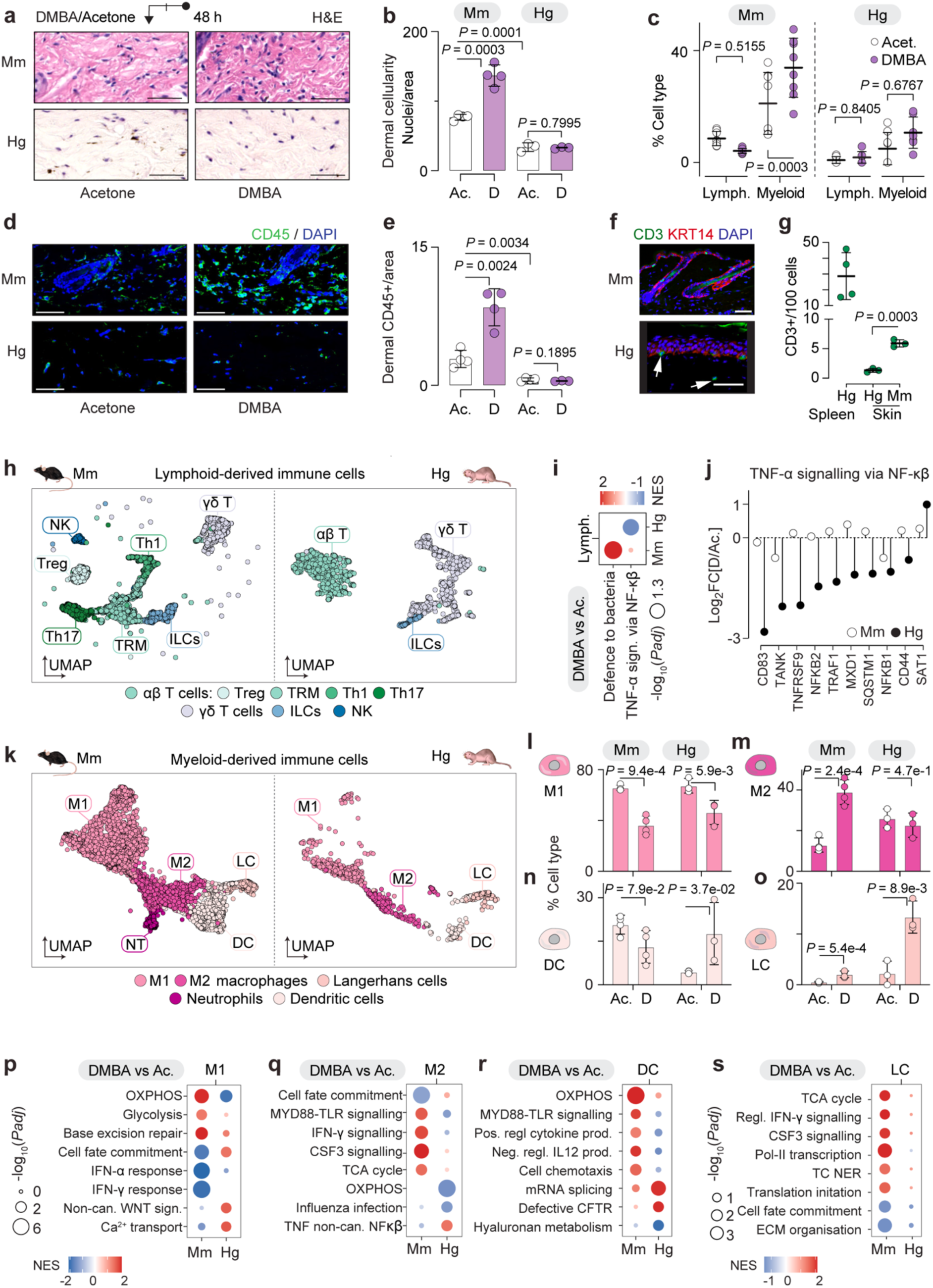
Restricted lymphoid and myeloid immune activation in naked mole-rat skin following acute carcinogen exposure. **a**, **b**, H&E-stained skin sections (**a**) showing mouse (*Mm*) and naked mole-rat (*Hg*) dermis 48 h after topical acetone (Ac.) or DMBA (D) application and quantification of dermal cellularity in *Mm* and *Hg* skin (**b**). **c**, Immune cell composition of the non-epithelial skin compartment showing lymphoid and myeloid populations in *Mm* and *Hg* skin treated with acetone or DMBA for 48 h. Data points show biopsies. **d**, **e**, Immunofluorescence (**d**) and quantification (**e**) of CD45⁺ immune cells in *Mm* and *Hg* dermis 48 h after Ac.- or D-treatment. **f**, **g**, CD3⁺ T cell immunofluorescence in homeostatic *Mm* and *Hg* skin (**f**) and quantification in *Hg* spleen and *Hg* and *Mm* skin (**g**). Arrows indicate CD3^+^ T cells. **h**, UMAP of lymphoid immune cells in *Mm* (left) and *Hg* (right) skin. Treg, regulatory T; TRM, tissue-resident memory T; Th1/Th17, T helper 1/17. ILCs, innate lymphoid cells; NK, natural killer cells. **i**, Gene set enrichment analysis of lymphoid-derived immune cells from DMBA-versus Ac-treated *Mm* and *Hg* skin. **j**, Log₂ fold change of leading-edge genes from the TNF-α signalling via NF-κB gene set in *Mm* and *Hg* skin after D-relative to Ac.-treatment. **k**, UMAP of myeloid-derived immune cells in *Mm* (left) and *Hg* (right) skin. M1, type 1 macrophages; M2, type 2 macrophages; DC, dendritic cells; LC, Langerhans cells; NT, neutrophils. **l–o**, Myeloid immune cell abundances in Ac.- or D-treated *Mm* and *Hg* skin. **p–s**, Gene set enrichment analysis of M1 macrophages (**p**), M2 macrophages (**q**), DCs (**r**) and LCs (**s**) comparing DMBA-versus Ac.-treated *Mm* and *Hg* skin. Circle size reflects significance; colour indicates NES. CFTR, cystic fibrosis transmembrane conductance regulator; TC-NER, transcription-coupled nucleotide excision repair. Scale bars, 50 μm (**a**, **d**, **f**). Values show mean ± s.d. (**b**, **e**, **g**, **l–o**) or median ± s.d. (**c**). N = 3–4 biologically independent individuals per species and treatment (**b**, **c**, **e**, **g–s**). Unpaired two-tailed *t*-test (**b**, **e**). Linear model (**c**, **l–o**). Exact *P* values are shown.

To test whether the sparsely distributed immune cells in naked mole-rat skin responded to the DMBA treatment, we first focused our studies on T cells as they play a key role in adaptive immunity by specifically recognizing pathogens, allergens, and tumor antigens ^43^. Both CD3 labeling and our integrated transcriptomic analysis in unperturbed skin revealed that naked mole-rats have very few T cells (2.1% ± 0.7) compared to human (9.9% ± 4.4) and mouse (6% ± 2.5) as previously reported ^22^ (Fig. 4f-g; Extended Data Fig. 6c-h). Higher-resolution clustering identified seven T cell subtypes in mice, but only three in naked mole-rats (Fig. 4h; Extended Data Fig. 7a-c), which is consistent with the simplified and less diverse T cell landscape recently reported in this species ^23,44^. Although the size of the T cell population remained unchanged following DMBA exposure (Extended Data Fig. 7c), they collectively showed reduced TNF signaling via NF-κB and a dampened pro-inflammatory response in naked mole-rats (Fig. 4i-j; Extended Data Fig. 7d-g). We concluded that naked mole-rat T cells were unlikely to play an active role in maintaining tissue integrity following DMBA exposure.

Then, we focused on myeloid-derived cells to explore potential compensatory innate immune responses. Cross-species comparison revealed two subpopulations of macrophages (M1 = regulatory/homeostatic macrophages; M2 = inflammatory/tissue-remodeling macrophages), neutrophils, Langerhans and dendritic cells (Fig. 4k; Extended Data Fig. 7h-k). DMBA treatment significantly reduced the fraction of M1 macrophages in both species, but only mice increased the population of M2 macrophages (Fig. 4l-m). In contrast, DMBA-treated naked mole-rat skin showed a significantly increased number of dendritic and Langerhans cells (Fig. 4n-o). However, unlike in mice, none of the naked mole-rat myeloid-derived immune cells transcriptionally upregulated inflammatory response pathways in response to DMBA (Fig. 4p-s; Extended Data Fig. 7l). Together our data showed that naked mole-rat immune cells prioritized regulatory programs maintaining tissue integrity and immune homeostasis over pro-inflammatory activation.

In summary, acute DMBA exposure in naked mole-rat skin elicits an immune response biased toward surveillance and tissue protection rather than inflammatory remodeling, consistent with a regenerative tissue response instead of increased cell death. However, the absence of cell loss in a highly regenerative environment raised the question of whether protection by an enlarged hybrid cell population, together with efficient genome-maintenance pathways, sufficiently prevented genotoxic lesions from persisting and propagating within the basal epidermal layer.

### Oncogenic stress leads to selective but constrained clonal expansion in naked mole-rats

Because sustained activation of DNA repair imposes a high metabolic demand, maintaining viability and genome integrity in non-proliferating cells becomes energetically costly over time ^45^. To test the limits of the tumor-suppressive capacity in naked mole-rats, we exposed the skin to chronic oncogenic stress and asked whether basal stem cells would be recruited to replenish the progenitor pool.

To examine this experimentally, we applied the two-stage chemical carcinogenesis protocol, in which a tumor-promotion phase follows DMBA exposure (Fig. 5a, top) ^46^. While all mice developed multiple epithelial skin tumors (papillomas) within 13 weeks, naked mole-rats remained tumor-free and exhibited no macroscopic, histological or cellular alterations of the epidermis, even after 24 weeks of treatment (Fig. 5a-c; Extended Data Fig. 8a-c). Nevertheless, treatment of skin with TPA resulted in enhanced proliferation and epidermal thickening in both species as expected (Extended Data Fig. 8d-f) ^47^. Under chronic oncogenic stress in naked mole-rats, basal stem cells were mobilized to replenish the arrested progenitor pool resulting in a significant reduction in K14/K10 double-positive cells within the basal layer and an increased stem and progenitor replacement rate (Fig. 5d-f).

**Fig. 5:**
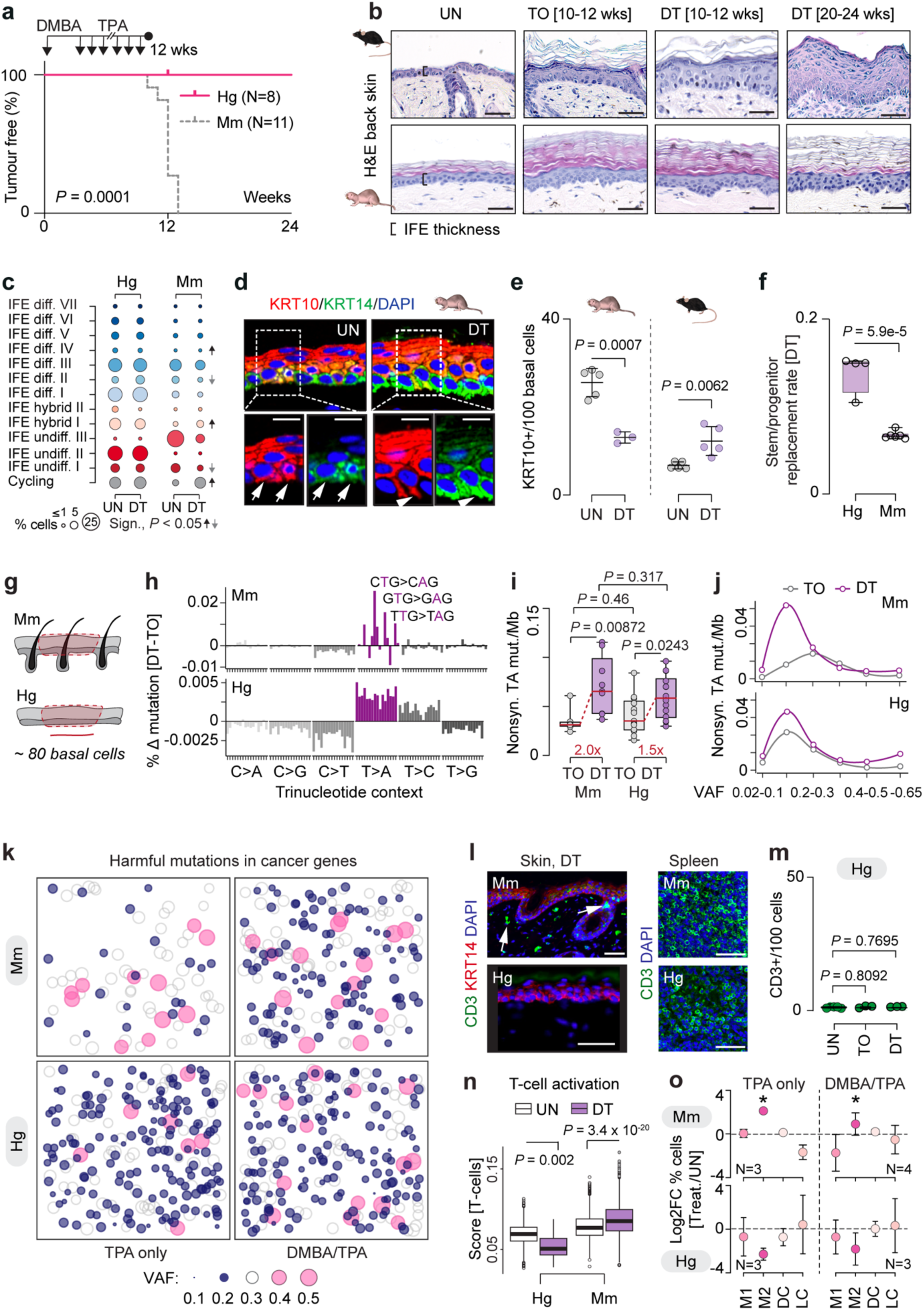
Naked mole-rats constrain expansion of mutant epidermal clones. **a**, Percentage of tumour-free animals following DMBA/TPA (DT) treatment of naked mole-rat (*Hg*) and mouse (*Mm*) back skin (*Hg*: 12 weeks, N = 4; 24 weeks, N = 4; *Mm*: 10–12 weeks, N = 11). **b**, H&E staining of untreated (UN), TPA-only (TO) or DT-treated (DT) back skin from *Mm* (top) and *Hg* (bottom) at the indicated time points. **c**, Proportion of IFE cell states in UN versus DT *Hg* and *Mm* skin. Significant (sign.) changes occur only in *Mm* (*P* < 0.05). *Hg*: UN, N = 5, n = 14; DT, N = 4, n = 11. *Mm*: UN, N = 3, n = 9; DT, N = 3, n = 5. **d**, **e**, Immunofluorescence (**d**) and quantification (**e**) of KRT14⁺KRT10⁺ basal cells (arrows) in UN and DT *Hg* and *Mm* skin; data points show means of ≥15 measurements per individual from 3–4 individuals. **f**, Stem and progenitor replacement rates after DT exposure inferred from clonal expansion; one point per biopsy (*Hg*, N = 4; *Mm*, N = 3). **g**, Schematic of laser-capture microdissection of IFE regions (∼80 basal cells). **h**, SNVs frequency across 96 trinucleotide contexts in DT-treated *Mm* (top) and *Hg* (bottom) IFE after subtraction of TO background. *Mm*: TO, N = 3, n = 2–3; DT, N = 3, n = 3. *Hg*: TO, N = 3, n = 4–5; DT, N = 4, n = 2–6. **i–k**, Nonsynonymous T>A mutation burden (**i**), mean burden by VAF bin (**j**), and nonsynonymous mutations in cancer and skin-cancer genes with predicted moderate or high functional impact (**k**), aggregated by treatment. Circle size and colour indicate VAF (≤0.2 or ≥0.4). Sample sizes as in **h**. **l**, **m**, CD3⁺ T-cell immunofluorescence in DT-treated *Mm* and *Hg* skin and UN spleen (l), with quantification in UN-, TO- and DT-treated *Hg* skin (m). Data points represent biological replicates (N = 3–5), averaged from ≥3 sections per individual. **n**, Gene set scores of *Hg* and *Mm* T cells in UN and DT skin. **o**, Log₂ fold change of myeloid immune proportions in TO- or DT-treated *Mm* and *Hg* skin relative to UN. *Mm*: TO, N = 3, n = 1; DT, N = 4, n = 1–5. *Hg*: TO, N = 3, n = 1; DT, N = 3, n = 1. Scale bars, 50 μm (**b**, **l**) and 10 μm (**d**). Box plots show quartiles and range (**f**, **i**) or 5th–95th percentile whiskers (**n**). Log-rank (Mantel-Cox) test (**a**). Linear model (**c**, **i**). Unpaired two-tailed *t*-test (**e**, **f**, **m**). Generalised linear model (**o**). \**P* < 0.05 (**o**). Exact *P* values are shown for all other panels.

This increased turnover raised the question of whether DMBA-initiated mutant clones are diluted and eliminated through progenitor cell-driven replacement, or instead persist and undergo clonal expansion within the interfollicular epidermis. To resolve this, we quantified mutational burden and clonal architecture following DMBA/TPA treatment in laser-captured micro-dissected epidermal regions encompassing ∼80 basal cells in both species (Fig. 5g; Extended Data Fig. 9a). This sampling scale maximized sensitivity for detecting rare somatic mutations and resolving clonal structure, while minimizing signal dilution from surrounding non-mutant cells ^48^. Whole-genome sequencing of the epidermal micro-biopsies revealed the expected DMBA-induced T>A transversions and a significant increase in non-synonymous T>A mutation burden in both species (Fig. 5h-i; Extended Data Fig. 9b-1).

Despite comparable mutational burdens, the clonal dynamics of epidermal cells carrying genotoxic lesions after DMBA/TPA treatment differed between the species. Mouse epidermis accumulated numerous low VAF non-synonymous T>A mutant clones (Fig. 5j; top panel), whereas the VAF distributions in naked mole-rat epidermis remained unchanged (Fig. 5j; bottom panel). This difference of TPA-driven clonal dynamics between the two species can be explained by the lower stem and progenitor replacement rate in the mouse epidermis (Fig. 5f). Initiated mutant clones persist and thereby create multiple independent opportunities for clonal outgrowth and malignant progression. In contrast, the higher base-line stem/progenitor turn-over in naked mole-rats outcompeted these new clones over time, even when these mutations occurred in cancer–associated genes (Fig. 5k). Notably, TPA treatment led to a higher number of clones with harmful mutations in naked mole-rats than mice, demonstrating that selective growth of rare mutant cells can occur even within an overall suppressive tissue environment (Fig. 5k; left panels).

Together, these findings support a model in which an expanded committed progenitor pool, coupled to elevated stem/progenitor turnover, efficiently dilutes DMBA-initiated mutant clones in naked mole-rat epidermis, thereby preserving long-term epidermal integrity (Fig. 5b-c; Extended Data Fig. 8c). This also explained the absence of overt immune activation following DMBA/TPA treatment in naked mole-rat skin. T cells did not accumulate and in addition, showed transcriptional repression of genes involved in their activation (Fig. 5l-n). Similar to short-term DMBA exposure, we detected no increase in M2 macrophages (Fig. 5o) and none of the myeloid cell types exhibited pro-inflammatory transcriptional signatures (Extended Data Fig. 9e-h). Collectively, our data showed that clonal expansion was selectively permitted in naked mole-rats but tightly constrained under chronic oncogenic stress.

### Hybrid basal cells are required to sustain cytostatic repair and immune quiescence

To directly test whether the K14⁺/K10⁺ hybrid population was functionally required for epidermal protection against oncogenic stress, we selectively depleted these cells from the basal layer. This was possible by first treating the skin with TPA, which reduced the number of hybrid cells in the basal layer of the epidermis by threefold (Fig. 6a). Then, we followed the classical two-step carcinogenesis DMBA/TPA protocol for 12 weeks (Fig. 6b). Morphologically, the TPA/DMBA/TPA treatment induced a more mouse-like stress phenotype, with significantly increased numbers of basal K14^+^/K10^+^ hybrid cells, more PCNA-positive cells, and greater epidermal thickening (Fig. 6c-h; Extended Data Fig. 10a). Thus, our data showed that the hybrid population was required to constrain epidermal responses under chronic carcinogenic challenge.

**Fig. 6:**
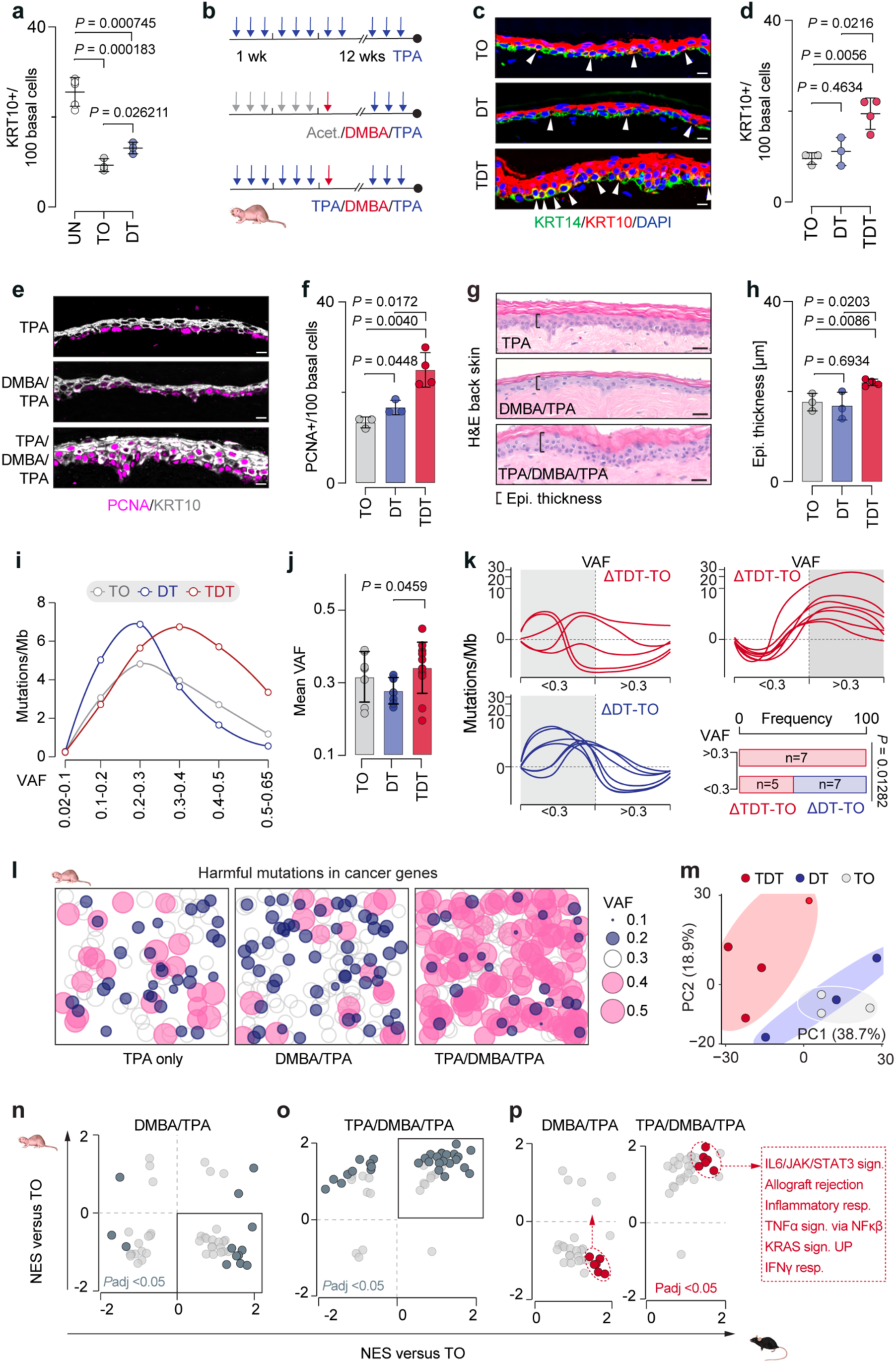
Disrupted progenitor cell balance sensitizes naked mole-rat skin to epithelial transformation. **a**, K10^+^ cells per 100 basal cells in back skin IFE of untreated (UN), TPA-only (TO), or DMBA/TPA (DT)-treated naked mole-rats (*Hg*) after 10–12 weeks. **b**, To perturb the K14⁺/K10⁺ hybrid compartment, *Hg* skin was treated with TPA for 2 weeks, followed by a single DMBA application and then TPA for 11 weeks. **c**, **d**, Immunofluorescence and quantification of K10^+^ cells. Arrowheads indicate K10^+^K14^+^ basal cells. **e**, **f**, Increased PCNA^+^ cells in TPA/DMBA/TPA (TDT)-treated *Hg* skin. **g**, **h**, H&E staining (**g**) and epidermal thickness quantification (**h**). **i**, **j**, SNV mutation burden by VAF in *Hg* IFE. **k**, Delta (Δ) SNV mutation burden versus VAF in *Hg* IFE, calculated as the difference between treatment and mean TPA controls per VAF bin; y axis uses inverse hyperbolic sine (asinh) transformation. Samples were classified by maximal Δ burden VAF (<0.3, ≥0.3). **l**, Nonsynonymous SNVs in cancer genes predicted to have moderate or high harmful impact in *Hg* IFE, aggregated by treatment. Each randomly plotted circle represents one mutated gene; size and colour indicate VAF. **m**, Principal component analysis of pseudo-bulk *Hg* IFE single-cell transcriptomes with confidence ellipses shaded by treatment. **n–p**, Gene set enrichment analysis of *Hg* IFE cells treated with DT vs TO (**n**, **p**; left) or TDT versus TO (**o**, **p**; right), contrasted with *Mm* IFE DT vs TO. Significantly enriched pathways (*P*adj < 0.05) are shown in dark grey (**n**, **o**); pathways gained in *Hg* TDT vs DT are shown in red (**p**). Scale bars, 10 μm (**c**, **e**). Values show mean ± s.d. (**a**, **d**, **f**, **h**). N = 3–6 biologically independent individuals per treatment (**a**, **b**, **d**, **h**). *Hg,* TO, N = 3, n = 2–3; DT, N = 3, n = 1–3; TDT, N = 4, n = 3 (**i–l**). *Hg,* N = 3-4 biologically independent individuals per treatment group (**m–p)**. Unpaired two-tailed *t*-test (**a**, **d**, **f**, **h**). One-way ANOVA (**j**). χ² test and Fisher’s exact test (**k**). Exact *P* values are shown.

To test how removal of the hybrid population affected the expansion of mutant clones, we micro-dissected epidermal regions (∼80 basal cells) from naked mole-rats exposed to the different treatment regimens and performed whole genome sequencing (Fig. 6b; Extended Data Fig. 10b-c). Consistent with our hypothesis that the hybrid population constrained clonal expansion under chronic oncogenic stress, we find a significant increase of clones with higher VAFs when we treated with TPA before DMBA exposure (Fig. 6i-j; Extended Data Fig. 10d-e). Seven out of 12 TPA/DMBA/TPA, but none of the DMBA/TPA micro-biopsies contained clones with VAFs greater than 0.3 (Fig. 6k), and this included harmful mutations in cancer genes (Fig. 6l). We concluded that the hybrid population was required for constraining the expansion of mutant clones when exposed to genotoxic stress.

To identify epidermal stress pathways that were activated upon chronic carcinogenic challenge in the absence of the hybrid population, we performed single cell transcriptome analyses. Epidermal cells exposed to TPA/DMBA/TPA exhibited a distinct gene expression programme when compared to cells treated with DMBA/TPA or TPA alone (Fig. 6m; Extended Data Fig. 10f-i). Notably these transcriptional changes were primarily driven by upregulation of inflammatory response pathways, identical to those activated by DMBA/TPA in the mouse epidermis (Fig. 6n-p). Thus, our data demonstrated that the hybrid progenitor population was an active part of the cellular mechanisms that mediated the naked mole-rat’s resistance to tumor initiation. This high turnover-driven cellular mechanism diluted mutagenic clones but permitted limited, selective clonal expansion within an otherwise strongly tumor-suppressive tissue environment.

### Dermal fibroblasts enforce a non-permissive niche for tumor progression

Clonal expansion of epithelial cells alone is insufficient for pre-cancerous progression and requires a permissive stromal environment ^49^. Cross-species integration of dermal fibroblast transcriptomes from naked mole-rat, human, and mouse skin identified multiple fibroblast subtypes per species (Fig. 7a, b; Extended Data Fig. 11a). Two states aligned between naked mole-rat and human, but not mouse, consistent with species- and site-specific dermal specialization and appendage-associated stromal programs. The species-specific fibroblast transcriptional states persist as stable, cell-intrinsic programs, because isolated dermal fibroblasts from mice and naked mole-rats have opposing effects on cancer cell growth. Only mouse fibroblasts promoted human squamous cell carcinoma (SCC25) proliferation in 2D and 3D *in vitro* co-culture models (Fig. 7c-e). We confirmed that naked mole-rat fibroblasts remained viable and proliferative under standard human cell-culture conditions (37 °C, 18% O_2_) (Extended Data Fig. 11b-i). Thus, naked mole-rat fibroblasts maintain a cell intrinsic stable, tumor-restrictive state that limits human SCC growth *in vitro*.

**Fig. 7:**
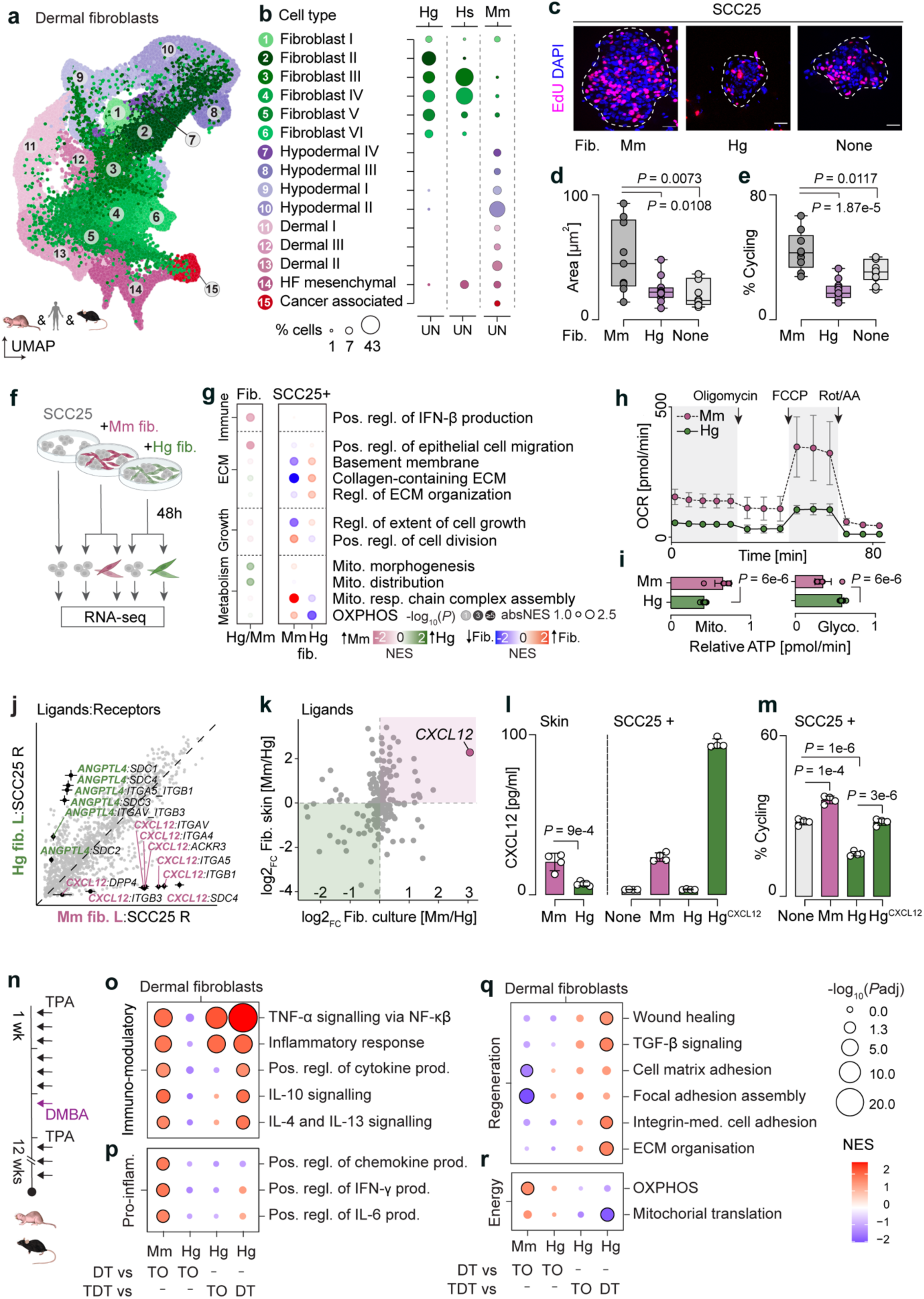
Naked mole-rat fibroblasts restrict pre-cancerous niches by creating a non-permissive microenvironment. **a**, **b**, UMAP and cell state assignment of skin fibroblasts from homeostatic naked mole-rat (*Hg*), human (*Hs*) and mouse (*Mm*) skin. UN, untreated; HF, hair follicle. *Hs*: N = 5, n = 5; *Hg*: N = 5, n = 14; *Mm*: N = 2, n = 2. **c–e**, EdU immunofluorescence (**c**) and quantification of EdU^+^ area (**d**) and % cycling cells (**e**) in human squamous carcinoma cells (SCC25) spheroids co-cultured with *Mm*, *Hg* or no fibroblasts. Experiments were performed independently three times; n = 9–11 spheroids per condition from one representative experiment. Scale bars, 10 μm (**c**). **f**, Schematic of 2D SCC25–fibroblast co-culture. **g**, Gene set enrichment analysis of orthologous differentially expressed genes in *Hg* versus *Mm* fibroblasts (**g**, left) or in SCC25 (**g**, right) co-cultures; n = 4–5 per species. Data points are shaded by significance, sized by absolute NES and coloured by direction of change. **h**, **i**, Oxygen consumption rate (**h**) and proportion of ATP production from mitochondrial respiration or glycolysis (**i**) in *Mm* and *Hg* fibroblasts. Experiments were performed independently three times; n = 8-9 technical replicates per condition from one representative experiment. **j**, Ligand–receptor expression correlation between SCC25 receptors and *Hg* or *Mm* fibroblast ligands. **k**, Comparative *Mm* versus *Hg* fibroblast ligand expression, co-cultured with SCC25 (x axis) vs in homeostatic skin (y axis). n = 4–5 per cell type (**j**, **k**). **l**, CXCL12 protein levels in *Mm* and *Hg* skin and in SCC25 co-culture conditions, including CXCL12-overexpressing *Hg* fibroblasts. **m**, EdU^+^ cycling SCC25 cells in 2D co-cultures. Skin samples: *Mm*, N = 4; *Hg*, N = 5 biologically independent individuals (**l**, left). Cell culture samples: n = 4 per condition (**l**, right; **m**). Experiments were performed independently three times; one representative experiment is shown (**l**, **m**). **n–r**, Schematic of the TPA/DMBA/TPA (TDT) carcinogenesis perturbation experiment (**n**) and gene set enrichment analysis of skin fibroblasts (**o–r**). N = 3-4 biologically independent individuals per treatment group (**o–r)**. Box plots show first quartile, median, third quartile and range (**d**, **e**). Values show mean ± s.d. (**h, i**, **l**, **m)**. Unpaired two-tailed *t*-test (**d**, **e**, **i**, **l**, **m**). Exact *P* values are shown.

To determine how this non-permissive stromal barrier to pre-cancerous progression was formed, we isolated fibroblast and tumor cells after 48 hours of co-culture and performed transcriptome profiling (Fig. 7f). Naked mole-rat fibroblasts resisted inflammatory activation but activated mitochondrial morphogenesis programs consistent with energetic stress, while imposing a metabolic brake and triggering ECM remodeling in SCC25 cells (Fig. 7g).

Metabolic plasticity is a key determinant of tumor-supportive fibroblast functions ^50^. Therefore, we asked whether the mitochondrial changes in naked mole-rat fibroblasts reflected a transient co-culture response or an intrinsic metabolic feature that limits their tumor-supportive capacity. To test this, we measured mitochondrial respiration and found that naked mole-rat fibroblasts had reduced basal and maximal oxygen consumption rates (OCR) with increased glycolysis, resulting in lower ATP production and decreased spare respiratory capacity relative to mouse fibroblasts (Fig. 7h-i; Extended Data Fig. 11j-n). Thus, an intrinsic metabolic constraint in naked mole-rat fibroblasts precludes tumor-supportive stromal functions.

Given that extrinsic stromal signals can reprogram metabolism ^51^, we used receptor-ligand analyses to pinpoint species-specific fibroblast-derived factors linked to metabolic plasticity. While most predicted fibroblast-SCC25 ligand-receptor interactions were conserved between mouse and naked mole-rat co-cultures, ANGPTL4 and CXCL12 emerged as species-specific signals (Fig. 7j; Extended Data Fig. 11o-p). Whereas ANGPTL4 can promote aerobic glycolysis ^52,53^, CXCL12 is sufficient to reprogram fibroblasts into a cancer-associated fibroblast (CAF)-like state ^54^, CAFs undergo metabolic reprogramming to supply energy-rich nutrients that fuel OXPHOS in adjacent cancer cells ^55^. Moreover, CXCL12 is a well-established epithelial mitogen and was consistently up-regulated in mouse but not naked mole-rat fibroblasts on both RNA and protein levels (Fig. 7k-m). In mouse fibroblasts, predicted CXCL12-receptor interactions were associated with activation of canonical pro-proliferative pathways, including JAK-STAT, MAPK/ERK and WNT/β-catenin signaling (Fig. 7m; Extended Data Fig. 11q). Forced CXCL12 expression in naked mole-rat fibroblasts was sufficient to restore their tumor-supportive activity (Fig. 7l–m). Accordingly, naked mole-rat fibroblasts also failed to support tumor growth in xenotransplantation assays into host mice (Extended Data Fig. 12a-d).

Finally, we asked how the tumor-restrictive, non-inflammatory fibroblast program, marked by absence of CXCL12, responded to DMBA in the absence of the K14⁺/K10⁺ hybrid population (Fig. 7n). Under these conditions, IL-10, IL-4 and IL-13 signaling was upregulated in fibroblasts of both species (Fig. 7o), whereas naked mole-rat fibroblasts failed to induce pro-inflammatory cytokine programs (Fig. 7p). This response was accompanied by selective upregulation of regenerative and tissue-repair–associated pathways, including wound-response genes, TGF-β signaling, and ECM organisation (Fig. 7q), together with an absence of mitochondrial remodeling in naked mole-rat fibroblasts (Fig. 7r). These transcriptional changes are consistent with a regulatory, tissue-protective stromal program rather than a tumor-permissive fibroblast state.

Collectively, our study shows that naked mole-rat skin sustains high mutational burden and rapid turnover through an expanded pool of differentiation-committed basal progenitors that preserves epidermal integrity via replication-coupled DNA repair. Immune quiescence and a tumor-restrictive fibroblast niche then favor clone dilution over progression.

## Discussion

Cancer progression is a multistep evolutionary process driven by accumulating genetic alterations that rewire cell identity and progressively remodel the tumor microenvironment ^49^. Different organs rely on distinct tumor-suppressive strategies, and skin is a particularly relevant tissue for understanding how proliferation, mutagenesis and immune activation intersect. As a rapidly renewing tissue, UV-exposed skin has a dense mosaic of competing clones, many of which harbor canonical cancer driver mutations, yet they persist within a largely intact tissue architecture ^56^. We uncover that naked mole-rat skin matches or exceeds mouse mutation rates, yet mutational burden and rapid turnover remain uncoupled from cancer risk, and even chronic carcinogenic exposure failed to drive malignant progression.

Continuous clonal replacement can dilute or extinguish initiated lineages unless they acquire a sufficient fitness advantage to persist and expand ^57^. *In vivo* lineage tracing shows that interfollicular epidermal homeostasis can be sustained by stochastic self-renewal and clonal competition consistent with neutral drift ^58^. Regulated fate decisions and commitment to differentiation can further limit proliferative potential and shorten the window for clonal selection ^59^. In naked mole-rat skin, we identify an unusually large population of committed K14⁺/K10⁺ hybrid basal cells, a rare state in mouse and human epidermis. This hybrid population is primed for terminal differentiation and has limited proliferative capacity. We propose that this expanded committed compartment buffers genotoxic and oncogenic stress by absorbing proliferative demand and constraining self-renewal, thereby restricting clonal expansion while preserving tissue integrity.

Tumor suppression can reflect tissue-level constraints, whereby niche competition and architecture shape clonal fate in addition to genetic change ^60^. In the epidermis, damaged cells can be eliminated through differentiation-linked delamination or cell death, followed by immune clearance ^61–63^. We find no evidence for increased cell death in naked mole-rat epidermis following genotoxic stress. Instead, our depletion experiments identify the K14⁺/K10⁺ hybrid basal population as an integral component of a protective epidermal program that preserves tissue integrity through regulated turnover and repair.

However, our data do not support a model in which tumor suppression arises from hybrid cell state alone. Naked mole-rat skin establishes a non-permissive stromal and immune microenvironment that constrains clonal expansion and malignant progression despite substantial mutational burden. We uncover that naked mole-rat dermal fibroblasts enforce a stable, non-permissive niche for tumor progression. Compared with mouse, naked mole-rat fibroblasts resist inflammatory/CAF-like activation and fail to induce tumor-supportive signals such as CXCL12.

Tumor-supportive CAFs can undergo energy-metabolic reprogramming to generate and export metabolites that fuel tumor bioenergetics and biosynthesis ^64^. In contrast, naked mole-rat fibroblasts are metabolically constrained showing reduced mitochondrial respiration and spare respiratory capacity with a shift toward glycolysis. Although tumor-supportive CAFs often become glycolytic, this typically reflects an activated, high-flux state that preserves energetic capacity and metabolic plasticity ^65^. Limited metabolic flexibility in naked mole-rat fibroblasts likely constrains energy-demanding tumor-supportive functions, such as sustained matrix remodeling and paracrine signaling, reinforcing a non-permissive stromal barrier to progression.

## Material and Methods

### Animals and tissue samples

All mice were housed in the DKFZ Central Animal Laboratory. Experimental procedures using C57BL/6J mice (Janvier) or NSG mice (Charles River) were performed under the terms and conditions of the experimental animal project licenses DKFZ365, G-59/17, G-96/21, G-215/19, and G-351/19, following approval by the local governmental ethics committee. Naked mole-rats were housed at the University of Duisburg-Essen and the University of Cambridge. Maintenance of naked mole-rats housed at the University of Duisburg-Essen was approved by the Veterinary Office of the City of Essen (AZ: 32-2-1180-71/328). Tissue sampling from these naked mole-rats was carried out in accordance with the German Regulations for Laboratory Animal Science (GV-SOLAS). Maintenance and skin carcinogenesis treatment of naked mole-rats housed at the University of Cambridge were performed under the terms and conditions of the UK Home Office license P7EBFC1B1. Fibroblasts were obtained from animals housed at the University of Cambridge, and experiments were carried out under Home Office license P7EBFC1B1. All animal experiments conducted in the UK were in accordance with the UK Animals (Scientific Procedures) Act 1986 Amendment Regulations 2012. BrdU/EdU cell-labeling experiments in naked mole-rats were approved by LANUV (Landesamt für Natur-, Umwelt- und Verbraucherschutz) NRW (81-02.04.2021.A472). For BrdU and EdU cell-labeling experiments, a conditional transgenic mouse line (K14CreER/SRSF2/tdTom) was used without induction by 4-hydroxytamoxifen (4-OHT). In the absence of 4-OHT, Cre recombinase is not activated, and these mice are therefore effectively wild type.

### Human skin samples

Formalin-fixed paraffin-embedded (FFPE) skin samples from five human donors were used for immunofluorescence. A scapular skin sample from a healthy 55-year-old male was obtained from Origene (PA15476B6E). Two abdominal skin samples were obtained from Genoskin: one from a 27-year-old male with normal pathology and Caucasian skin type III (20191105.07), and one from a 46-year-old male with normal pathology and Caucasian skin type III (20200213.09). Two additional axillary skin specimens were obtained from remnant tissue, not required for diagnostic purposes, of patients undergoing routine surgery at the Department of Dermatology, University Hospital of Heidelberg, after written informed consent from each patient and as approved by the Ethics Committee of Heidelberg University (S-091/2011). In both cases, skin with normal histopathology, collected at sites distant from any skin pathology, was provided: one specimen from a 48-year-old male and one from a 64-year-old male, neither of whom had evidence of chronic cutaneous photodamage or systemic immunosuppression.

### Chemical skin carcinogenesis

For short-term mutagen exposure, female mice (C57BL/6J, 7 weeks old) received a single topical application of 500 µg 7,12-dimethylbenz[a]anthracene (DMBA; Sigma-Aldrich) dissolved in 200 µl acetone or acetone vehicle alone and were euthanized 48 h later. Naked mole-rats (Essen colony, 2.5–3 years old; three females and one male per group) were treated with 500 µg DMBA in 250 µl acetone to account for their larger dorsal surface area.

For long-term carcinogenesis experiments, female mice (C57BL/6J) and male naked mole-rats were treated once with DMBA, followed by repeated applications of 12-O-tetradecanoylphorbol-13-acetate (TPA; Sigma-Aldrich) as previously described ^46^. Mice received 100 µg DMBA in 200 µl acetone, followed one week later by 10 µg TPA in 100 µl acetone, applied three times per week for up to 20 weeks. Male naked mole-rats (Cambridge colony) received 500 µg DMBA in 250 µl acetone, followed one week later by 60 µg TPA in 250 µl acetone, applied three times per week for up to 24 weeks. Control groups were treated with TPA alone (same concentration and schedule) for 10-12 weeks.

For the TPA-induced perturbation experiment, naked mole-rats (Essen colony, 4.5 years old) were divided into three groups: (1) TPA only, receiving 60 µg TPA in 250 µl acetone three times per week for 13 weeks; (2) acetone vehicle for two weeks, followed by a single DMBA dose (500 µg in 250 µl acetone) and subsequent TPA treatment, applied three times per week for up to 13 weeks; and (3) TPA treatment three times a week for two weeks, followed by a single DMBA dose and continued TPA treatment, applied three times per week for up to 13 weeks. Both sexes were included in all experimental groups. Animals used in this study are listed in Extended Data Table 1.

### Single cell isolation

For detailed information on 10x scRNA-sequencing samples, see Extended Data Table 2. Biopsies (1-5 mm³) from naked mole-rat or shaved mouse back skin were incubated using one of the following protocols: (i) dissociation buffer (1.25 mg/ml collagenase I (Sigma), 0.5 mg/ml collagenase II (Worthington), 0.5 mg/ml collagenase IV (Worthington), and 0.1 mg/ml hyaluronic acid (Sigma) in Hanks’ Balanced Salt Solution (HBSS; Invitrogen)) for 2.5 h, followed by addition of trypsin (Santa Cruz Biotechnology) to a final concentration of 0.25% for 30 min; or (ii) 0.25% trypsin (Santa Cruz Biotechnology) for 2 h, followed by incubation with dissociation buffer for 1 h. All dissociation steps were performed at 37 °C with shaking at 600 rpm. After 3 h, enzymatic digestion was quenched with HBSS supplemented with 10% fetal bovine serum (FBS; Life Technologies), and cell suspensions were filtered through 40 µm cell strainers (Greiner Bio-One). Strainers were washed with an additional 2–3 ml HBSS containing 10% FBS, and cells were pelleted by centrifugation at 300 × g for 8 min at 4 °C. Cell pellets were carefully resuspended (using wide-bore P1000 pipette tips) in 100–200 µl sorting buffer (1 mM EDTA (Invitrogen) and 0.04% HyClone bovine serum albumin (BSA; GE Healthcare Life Sciences) in PBS). Cell suspensions were either passed through a 40 µm FLOWMI cell strainer (Fisher Scientific), and viability and concentration were determined using 0.4% trypan blue stain (Invitrogen) and a Countess II automated cell counter (Invitrogen), or were subjected to fluorescence-activated cell sorting (FACS).

For FACS, cell pellets were resuspended in sorting buffer containing 1 µg/ml propidium iodide (PI; Life Technologies) and transferred to low-binding polypropylene FACS tubes. Live cells from each sample were sorted into a single well of a 96-well flat-bottom, low-attachment tissue culture plate containing 30 µl sorting buffer on a cold plate, using an Aria II flow cytometry cell sorter (BD Biosciences) with a 70 µm nozzle (70 psi). PI was excited with a 561 nm laser, and fluorescence was recorded using a 610/20 bandpass filter with a 595 nm long-pass filter. Cells were gated as follows: (i) FSC-A vs. SSC-A to remove debris; (ii) SSC-W vs. SSC-A to exclude doublets; and (iii) RL780/60-A vs. PI-A to exclude autofluorescent and dead (PI-positive) cells. Samples were handled on ice throughout.

### Single-cell capturing, library preparation and sequencing

10x Genomics Chromium Single Cell 3′ cDNA libraries were prepared according to the manufacturer’s instructions. For mouse samples, all sorted cells from a single 1 mm³ biopsy (800-1,100 untreated cells and 2,900-5,300 DMBA/TPA-treated cells) were loaded onto the 10x Chromium Controller. For all other 5 mm³ biopsy samples (from both mouse and naked mole-rat), cell suspensions were diluted to a concentration targeting recovery of 5,000 cells. Briefly, single cells were encapsulated into droplets in the Chromium Controller for cell lysis and barcoded reverse transcription (RT) of mRNA, followed by amplification, shearing, and Illumina library construction. Library quantification was performed using the Qubit dsDNA HS Assay Kit (Life Technologies), and cDNA integrity was assessed using D1000 ScreenTapes (Agilent Technologies). Libraries were pooled by species, and each pool was sequenced on an Illumina NextSeq 2000, NovaSeq 6000, or NovaSeq X Plus using either paired-end sequencing with 28 bp for read 1 and 94 bp for read 2, or 100 bp for each read.

### Histology, immunohistology and immunofluorescence staining

Tissues or tumors were fixed overnight with 4% paraformaldehyde (PFA) (Santa Cruz Biotechnology), transferred to 70% ethanol and then embedded into paraffin. Paraffin samples were cut into 4-5 μm (H&E) or 5-20 μm (RNAscope and immunofluorescence) sections. Paraffin sections were deparaffinized in xylene and rehydrated in a gradient of ethanol and distilled water and stained with haematoxylin and eosin (H&E) according to standard protocols.

For immunohistochemistry, tissue sections were deparaffinized in xylene, rehydrated through a graded ethanol series, and subjected to heat-induced epitope retrieval (pH 6.0, Target Retrieval Solution, Agilent Technologies). Sections were incubated for 1 h with the primary antibody against phospho-histone H2A.X (Ser139) (20E3, rabbit monoclonal, 1:100; #9718, Cell Signaling Technology). Positive signal was visualised using an AP-conjugated anti-rabbit secondary antibody (Signal Boost, Cell Signaling Technology) and a Vibrant Red Cell substrate (Cell Signaling Technology).

For immunofluorescent labelling of paraffin sections, sections were deparaffinized in xylene and rehydrated in a gradient of ethanol and distilled water and epitopes were retrieved by incubating slides for 20 minutes at 95 °C in 10 mM citrate buffer (pH 6.0) with 0.05% Tween-20. Sections were then blocked for 1 hour with 10% FBS/0.1% Triton-X at room temperature (RT). Primary antibodies were diluted in 1% FBS and incubated overnight using the following dilutions: mouse monoclonal KRT14 (1:1000, ab7800, Abcam), rabbit monoclonal KRT10 (1:200, ab76318, Abcam), mouse monoclonal KRT6 (1:100, LHK6B, ThermoFisher Scientific), rabbit polyclonal CD3 (1:200, ab5690, Abcam), rabbit polyclonal Ki67/MKI67 (1:200, ab15580, Abcam), rabbit monoclonal Ki67/MKI67 (1:100, ab16667, Abcam), rat monoclonal BrdU (1:100, ab6326, Abcam), rabbit polyclonal CD45 (1:200, ab10558, Abcam) and mouse monoclonal PCNA (1:2000, ab29, Abcam). Fluorescence-labelled Alexa Fluor-647, -555 and 488-conjugated secondary antibodies (Thermo Fisher Scientific) were added at a dilution of 1:1000 for 1 hour at RT to detect primary antibodies. DAPI 1:1000-3000 (Sigma Aldrich) was used as nuclear counterstain. Slides were mounted using fluorescent mounting media (Agilent). Antibody specificity in naked mole-rat tissues was validated using western blot, RNAscope or immunofluorescence (Extended Data Fig. 12e-j).

To visualize EdU labelled cells in mouse and naked-mole-rat skin tissues, paraffin embedded sections underwent staining following the immunofluorescence procedure described above, with additional steps: After epitope retrieval, sections were permeabilized using 0.1% Triton-X-100 in TBS for 10 minutes at RT. DNA denaturation was achieved by treating the slides with 2N HCl for 30 minutes at 37°C. Following a PBS wash, slides were exposed to 0.1% saponin diluted in PBS for 5 minutes at RT. To conjugate EdU with a fluorochrome, a 30-minute azide-alkyne cycloaddition was performed in the dark at room temperature using the Click-iT Cell Reaction Buffer Kit (Life Technologies) and 5 μM Alexa Fluor 488-Azide (Life Technologies). Sections were washed twice with 0.1% saponin/PBS buffer and blocked as described above.

### TUNEL assay

Apoptotic DNA fragmentation was detected using the TUNEL Assay Kit (ab66110; Abcam) with minor modifications. Formalin-fixed paraffin-embedded (FFPE) skin and spleen sections were deparaffinized in xylene (2 × 5 min), rehydrated through a graded ethanol series followed by 0.85% NaCl, and rinsed in PBS. Sections were permeabilized with Proteinase K (10 min), with DNase I (Qiagen, #79254; 15 min, 37 °C) applied to positive controls. After PBS washes, tissues were post-fixed in 4% paraformaldehyde (5 min) and equilibrated in PBS. DNA strand breaks were labelled with DNA Labelling Solution (TdT reaction buffer, TdT enzyme, Br-dUTP) for 1.5 h at 37 °C in a humidified chamber, followed by PBS washes and 30 min incubation with anti-BrdU-Red antibody at room temperature in the dark. Negative controls were processed in parallel without TdT enzyme. Nuclei were counterstained with DAPI (1:3000), rinsed in ddH₂O, and mounted in aqueous fluorescent medium (Agilent Technologies).

### Western blot

Proteins were extracted from naked mole-rat or mouse tissues using a polytron homogenizer in ice-cold RIPA buffer (50 mM Tris-HCl, pH 7.4, 1% NP-40, 150 mM NaCl, 0.1% SDS, and 0.5% sodium deoxycholate), supplemented with complete protease inhibitor (Roche). Protein concentrations were determined using the Pierce BCA Protein Assay kit (Thermo Fisher Scientific) and measured with a spectrophotometer. For each sample, 30 μg of protein lysates were mixed with Laemmli buffer (2X), separated on polyacrylamide gels (Bio-Rad), and blotted onto PVDF membranes (GE Healthcare). After blocking for 1 h at room temperature in 5% non-fat milk in TBST buffer (1X TBS and 0.1% Tween-20), the membranes were incubated overnight at 4 °C with KRT14 (1:1000, ab7800, Abcam), KRT10 (1:100, ab76318, Abcam), or KRT6 (1:1000, LHK6B, ThermoFisher Scientific) in blocking buffer. The following day, each membrane was washed three times for 10 min in TBST before incubation with either anti-mouse or anti-rabbit horseradish peroxidase (HRP)-conjugated secondary antibody (1:10,000, BioVision) in TBST at room temperature for 1 h. Finally, the membranes were washed four times with TBST, and bands were detected using the Clarity ECL detection reagent (Bio-Rad) with the ChemiDoc MP Imaging System (Bio-Rad).

### RNA in situ hybridisation

FFPE blocks of mouse and naked mole-rat skin and spleen fixed in 4% paraformaldehyde and cut into 5 µm (spleen) or 20 µm (skin) sections, were subjected to RNA FISH using the RNAScope Multiplex Fluorescent Detection Kit V2 Assay (ACDBio, cat. no. 323100) with TSA Vivid Fluorophores (Bio-Techne) according to the manufacturers’ instructions. Sample pretreatment steps included a 10 min incubation with hydrogen peroxide, a mild target retrieval at 98–102 °C for 15 min and incubation with RNAscope protease plus for 15 min at 40°C. Target probes (*Mm-Cd3d*: 460101-C1; *Mm-Krt10*: 457901-C2; *Mm-Krt14*: 422521-C3; *Hg-Cd3d*: 1146881-C1; *Hg-Krt10*: 1146891-C2; *Hg-Krt5*: 1168011-C3) were hybridized to the pre-treated sections for 2 hours at 40°C. Sections were counterstained with DAPI (1:5000, Sigma-Aldrich) or WGA (1:50, Invitrogen) and mounted using ProLong Gold Antifade Mountant (ThermoFisher).

### Cell culture, transfection and infection

Human squamous carcinoma cells (SCC25) (CTR-1628) and immortalised 3T3 mouse embryonic fibroblasts (MEFs) (CRL-2752) were obtained from ATCC. SCC25 cells stably expressing GFP and luciferase (SCC25-Luc-GFP) were kindly provided by S.A. Benitah ^66^. Transformed naked mole-rat fibroblasts were generated as described in Hadi et al., 2020 ^9^.

Unless otherwise stated, MEFs and SCC25 were cultured in a humidified 37°C incubator with 5% CO_2_ and 18% O_2_, while naked mole-rat fibroblasts were grown in a humidified 32°C incubator with 5% CO_2_ and 3% O_2_. SCC25 cells were cultured in DMEM/F12 medium (11320-032, Life Technologies) supplemented with 10% FBS (Life Technologies), 1% Penicillin-Streptomycin (PEST) (Sigma-Aldrich), 400 ng/ml hydrocortisone (Santa Cruz Biotechnology) and 1% Primocin Ant-pm-2 (InVivogen). MEFs and naked mole-rat dermal fibroblasts were cultured in DMEM high glucose (41965-039, Life Technologies) supplemented with 15% FBS (Life Technologies), 1% PEST (Sigma-Aldrich), 1% Sodium pyruvate (Life Technologies), 1% non-essential amino acids (Life Technologies) and 1% Primocin (Ant-pm-2, InVivogen).

To fluorescently label cells, Lenti-X 293T cells were grown in DMEM (Life Technologies) supplemented with 10% FBS (Life Technologies), 1% penicillin-streptomycin (Life Technologies). The cells were then transfected with either LeGO-iV2-Luc2-mVenus2, pLenti-C-mGFP-P2A-Puro, or pCDH-EF1-Luc2-P2A-tdTomato (Addgene) plasmids, along with packaging vectors, using TurboFect as per the manufacturer’s instructions (Life Technologies). Fibroblasts or SCC25 cells were subjected to two rounds of transduction using 0.45 µm filtered media containing virus particles and supplemented with 8 µg/ml polybrene (Sigma Aldrich) to enhance transduction efficiency. Twenty-four hours after transduction, the medium was replaced with respective cell culture medium, and the cells were expanded before FACS sorting.

For two-dimensional (2D) co-culture, SCC25-tdTomato cells were seeded in a 1:1 ratio with mouse or naked mole-rat fibroblasts in DMEM/F12 medium (11320-032, Life Technologies) supplemented with 0.05% FBS (Life Technologies), 400 ng/ml hydrocortisone (Santa Cruz Biotechnology) and 1% Penicillin-Streptomycin (PEST) and cultured in a 37°C incubator with 5% CO_2_. After 48 hours SCC25-tdTomato and fibroblasts were separated using FACS sorting.

To test the effect of forced CXCL12 expression in 2D co-culture conditions, naked mole-rat fibroblasts were seeded in at sub-confluency and transiently transfected the following day with a pcDNA3-CXCL12-sfGFP (Addgene, #98961) expression plasmid using Lipofectamine 2000 (Invitrogen) according to manufacturer’s instructions. Cells were maintained at 32 °C and 3% O₂. Twenty-four hours post-transfection, cells were seeded for 2D co-culture with SCC25 cells in F12/DMEM containing 0.5% FBS. Co-cultures were maintained at 37 °C and 18% O₂ for 48 hours.

For 3D sphere co-culture, SCC25 cells were mixed with mouse or naked mole-rat fibroblasts in a 1:1 ratio in SCC25 media, plated in ultra-low adherent culture dishes (Fisher Scientific) and cultured in a humified 37°C incubator with 5% CO_2_ for up to 6 days.

All cells routinely tested negative for mycoplasma infection.

### Flow cytometry and analysis

To separate co-cultured SCC25-tdTomato cells and fibroblasts from mice or naked mole-rat fibroblasts for RNA-sequencing, the cells were trypsinized, washed twice with ice-cold PBS containing 10% FBS (Life Technologies) and 1% sodium azide (Sigma-Aldrich). Subsequently, the cells were incubated with APC-conjugated E-Cadherin antibody (1:200, 130-111-840, Miltenyi Biotec or 1:00, 50-3249.82, Invitrogen) in ice-cold PBS containing 3% BSA (GE Healthcare Life Sciences). After a 30-minute incubation at 4 °C, the cells were washed twice in PBS. SCC25 cells were then sorted as tdTomato- and E-cadherin-positive, while fibroblasts were sorted as tdTomato- and E-cadherin-negative cells. Data acquisition was carried out using a cell sorter (BD Biosciences) and analysed using FlowJo (version 10.8.2).

### ELISA

To quantify CXCL12 protein levels in two-dimensional co-cultures, conditioned media were collected 72 h post-transfection (48 h of co-culture), cleared by centrifugation (2,000 × g, 10 min), aliquoted, and stored at −20 °C. For skin tissue measurements, snap-frozen samples were homogenized in chilled 1X Cell Extraction Buffer PTR supplemented with Halt protease and phosphatase inhibitors (Thermo Scientific), incubated on ice for 20 min, and centrifuged at 18,000 × g for 20 min at 4 °C. Supernatants were collected, protein concentrations were determined by Pierce BCA protein assay (Life Technologies), and samples were analysed immediately or stored at −80 °C. CXCL12 levels were measured using mouse (ab100741) or human (ab100637) SDF1α/CXCL12 ELISA kits (Abcam) according to the manufacturers’ instructions. Comparable results were obtained with both kits (data not shown). CXCL12 is highly conserved across species, with 93% amino acid identity between human and mouse, 94% between human and naked mole-rat, and 95% between mouse and naked mole-rat (data not shown).

### Proliferation rate

Proliferation was assessed by EdU incorporation in fibroblast monocultures, two-dimensional SCC25–fibroblast co-cultures, two-dimensional mouse primary keratinocyte–naked mole-rat fibroblast co-cultures, and three-dimensional SCC25–fibroblast spheroid co-cultures. Mouse or naked mole-rat fibroblasts were cultured alone or co-cultured with SCC25 cells or mouse primary keratinocytes as indicated. Cells were incubated with 10 μM 5-ethynyl-2′-deoxyuridine (EdU, Thermo Fisher Scientific) for 1–1.5 h, with samples lacking EdU serving as negative controls. Fibroblast monocultures and 2D co-cultures were harvested by trypsinization, whereas 3D spheroids were collected by gentle centrifugation. Samples were washed with PBS, fixed in 1% paraformaldehyde for 15 min on ice in the dark, and permeabilized in PBS supplemented with 3% FBS and 0.1% saponin for 5 min at room temperature. EdU incorporation was detected by azide–alkyne cycloaddition using the Click-iT Cell Reaction Buffer Kit (Life Technologies) with 5 μM Alexa Fluor 488– or 647–azide for 30 min at room temperature in the dark. For analysis of proliferation in 2D co-cultures, SCC25 cells and mouse primary keratinocytes were distinguished from fibroblasts by surface staining with PE/Cy7-conjugated anti–E-cadherin (CD324, clone 67A4; BioLegend, 1:100). Prior to antibody staining, cells were blocked in PBS containing 10% FBS and 1% sodium azide. Samples processed without antibody served as unstained controls. Samples were counterstained with DAPI (1 μg/ml). Single-cell suspensions were analyzed by flow cytometry (BD LSRFortessa or FACSCanto), while 3D spheroids were imaged using a TCS SP5 II confocal microscope (Leica).

### Cell viability

To assess the viability of fibroblasts in both 2D and spheroid co-cultures, cells were trypsinized, and cell suspensions were treated with 0.1 μg/ml DAPI for 15 minutes at room temperature. After a single wash with PBS, cells were directly analysed for the percentage of viable cells (DAPI-negative) or dead cells (DAPI-positive) using an LSRFortessa analyzer (BD Biosciences). A positive control sample for dead cells was included to establish gating parameters for identifying dead cells. To distinguish fibroblasts from SCC25 cells within the spheroids, mouse fibroblasts stably expressing GFP or naked mole-rat cells stably expressing mVenus were used.

### DMBA mutagenesis in fibroblasts

Sub-confluent (50%) MEFs and naked mole-rat fibroblasts were treated with 50 μM DMBA (D3254, Sigma-Aldrich) or vehicle control (0.05% DMSO) for 3 hours. Cells were washed and single sorted into 96 well plates using a 100 mm nozzle on an Aria I flow cytometry cell sorter (BD Biosciences). Strict gating was applied to exclude cellular debris and doublets using; i) SSC-A versus FSC-A, ii) SSC-W versus SSC-A and iii) FSC-H versus FSC-A. Single-sorted cells were manually inspected 3-24 hours after plating to exclude wells with more than one cell. MEF clones were expanded for a total of 14 days and naked mole-rat clones for 27 days. Genomic DNA was isolated from 200,000 cells from each expanded clone using the AllPrep micro DNA/RNA kit (Qiagen) according to the manufacturer’s instructions. gDNA quantity was determined using the Qubit dsDNA HS Assay Kit (Thermo Fisher) and gDNA integrity was assessed using a Genomic DNA ScreenTape (Agilent Technologies). The DIN was above 7.1 for all samples. Whole-genome sequencing (WGS) libraries were generated from 500 ng gDNA using the NEBNext Ultra II FS DNA Library Prep kit (NEB) following the manufacturer’s instructions. Library fragment size was determined using D1000 ScreenTapes (Agilent Technologies) and quantity determined using Qubit dsDNA HS Assay Kit (Thermo Fisher). Libraries were pooled in 6-plex per species and sequenced over three lanes HiSeq X (Illumina) lanes to produce paired-end 150-bp reads with a minimum 10X coverage. Deeper re-sequencing was done on NovaSeq X, paired-end 150-bp.

### Mitochondria assay

To quantify the number and activity of mitochondria, fibroblasts were treated with MitoTracker Deep Red and CMXRos Red (200 nM, Thermo Fisher Scientific) for 30 minutes at 37 °C. Following incubation, cells underwent two washes in PBS and were subsequently trypsinized. The fluorescence of each sample was then assessed using the BD LSRFortessa Analyzer (BD Biosciences). Fluorescence measurements were presented as histograms, and the fluorescence median values were used for quantification.

### Oxygen consumption, lactate production and ATP rate assay

Oxygen consumption rate (OCR) and extracellular acidification rate (ECAR) were assessed using a Seahorse XFe96 extracellular flux analyser (Agilent) with the Seahorse XF Cell Mito Stress Test Kit (Agilent), following the manufacturer’s guidelines. Briefly, cells were seeded at 3 × 10^4^ cells per well in cell culture microplates (Sigma-Aldrich) six hours before the assay to ensure equal cell attachment and to account for potential differences in cell number due to varying proliferation rates. This seeding process took place in a humidified 37°C incubator with 5% CO_2_ and 18% O_2_. After a one-hour equilibration period in XF assay medium supplemented with 10 mM glucose, 1 mM sodium pyruvate, and 2 mM glutamine in a 37°C incubator with 0% CO_2_, OCR and ECAR were monitored. Measurements were taken at baseline and during sequential injections of oligomycin (1 μM), carbonyl cyanide-4-(trifluoromethoxy)phenylhydrazone (1 μM), and rotenone or antimycin A (0.5 μM). Data for each well were normalised to protein concentration, determined using the Pierce BCA Protein Assay kit (Thermo Fisher Scientific). ATP production rate and spare respiratory capacity (SCR) was calculated according to manufacturer’s instructions.

### Imaging, processing and image analysis

Bright field images were acquired with either a Zeiss Axio Imager M2 Microscope and an Axiocam 506 colour camera or Zeiss Axioscan 7 using Zen (version 2.6) software (Carl Zeiss). Fluorescence microscopy images were acquired with either LSM 700 (Carl Zeiss) confocal microscope using Zen software, Zeiss Axioscan7 using Zen (version 2.6) software (Carl Zeiss) or TCS SP5 II (Leica) confocal microscope. All images were further processed using Fiji software (version 2.1.0). All measurements were performed either blinded for condition or by semi-automated macros.

### In vivo cell labelling

To measure the amount of cycling progenitor and quiescent long-term DNA-label-retaining stem cells (LRCs), cells were labelled using two thymidine analogues, BrdU and EdU ^29^. Actively growing mice (10 days old) and naked mole-rats (5 months old) were intraperitoneally injected four times with 50 mg/kg BrdU (Merck Millipore) in 12-hour intervals. 20 mg/kg of EdU (Biomol) was intraperitoneally injected one hour before each timepoint was collected. For three out of four naked mole-rats, two 3 mm^2^ punch biopsies per animal were collected four days after the initial BrdU injection. During this procedure, animals were anesthetized using ketamine (120 mg/kg) and xylazine (16 mg/kg) administered intraperitoneally. To alleviate postoperative pain, animals were provided with carprofen (up to 25 mg/kg). For LRC assays animals were chased for 10 weeks.

### Co-injection xenograft model

To investigate the influence of fibroblasts on tumor growth, we mixed one million human SCC25-GFP-Luc cells with: i) one million mouse fibroblasts (MEFs), ii) one million naked mole-rat fibroblasts, or iii) no fibroblasts. For assessing fibroblast growth, one million SCC25 cells were combined with: i) one million mouse fibroblasts expressing mVenus-Luc or ii) one million naked mole-rat fibroblasts expressing mVenus-Luc. Viability of all cells exceeded 95%. Cell mixtures were diluted in PBS and kept on ice until injection. A 100 µl volume of cell mixtures was subcutaneously injected into both flanks of 8-week-old female immunosuppressed NSG mice (*n* = 5-6 per experimental group) using a 22-gauge needle. Mice were anaesthetised with isoflurane gas during the injection procedure. Tumor growth was monitored for 28 days by measuring the luciferase bioluminescent signal (BLI) with the IVIS Lumina III In Vivo Imaging System (Perkin Elmer). For this, mice received an intraperitoneal injection of 50 μl 30 mg/ml d-luciferin (in PBS) (E1603, Promega) 5 minutes before imaging. Isoflurane gas was continuously administered to maintain anaesthesia during imaging. Data were quantified using Living Image software version 4.7 (Perkin Elmer). Regions of interest were manually outlined, and BLI signal was quantified as total photon flux (photons/second).

### DNA isolation from skin biopsies and livers

DNA was isolated from 1 mm^3^ mouse and naked mole-rat back skin biopsies and livers using a modified and combined version of the AllPrep DNA/RNA Micro Kit (Qiagen) and DNeasy Blood and Tissue Kit (Qiagen). Briefly, 350 μl RLT buffer (from the AllPrep DNA/RNA Micro Kit) containing 1% β-Mercaptoethanol (Sigma-Aldrich) was added to each sample (containing one biopsy) and the tissue was homogenised in a Qiagen TissueLyzer II using 2 cycles of 3 minutes at 30 Hz. After lysis 688 μl of nuclease-free water and 12 μl proteinase K (from DNeasy Blood and Tissue Kit) was added to the lysate and incubated at 56°C for 10 minutes. Half of the sample (519 ul) was used for DNA extraction. 519 μl Buffer AL (Qiagen Blood & Tissue) was added, mixed by vortexing, and incubated at 56°C for 30 minutes at 500 rpm. An equal volume (519 μl) of absolute ethanol was added to samples and mixed thoroughly. The precipitated mixture was transferred to a DNeasy Mini spin 96-well spin plate (Qiagen), washed twice with 700 μl buffer AW1 and twice with buffer AW2. After drying the membrane for 3 minutes at 20,000 x g DNA was eluted using two consecutive elutions of 30 µl pre-warmed AE buffer (70°C). The quality of the DNA was assessed using a Genomic DNA ScreenTape (Agilent Technologies).

### Whole-exome sequencing

Dual indexed whole exome capture libraries were prepared from mouse and naked mole-rat skin biopsies and matched livers using SureSelect XT HS and XT Low Input Enzymatic Fragmentation protocol (Agilent). For mouse whole exome sequencing (WES) libraries SureSelectXT Low Input Target Enrichment System was used with Mouse All Exon capture kit (49.6 Mb). For naked mole-rat WES libraries SureSelectXT HS / SureSelectXT Low Input Target Enrichment with Pre-Capture Pooling was used with a custom-designed NMR all exon capture array (Agilent). Exome libraries were sequenced on Illumina NextSeq2000 and NovaSeq6000 sequencers using a 100 bp paired-end read protocol.

### Bulk RNA-sequencing

To analyse the transcriptomes from the 2D co-cultures, approximately 50,000-100,000 cells of each cell type (SCC25 or fibroblasts) from each condition were sorted. Total RNA was extracted by adding 800 µl TRIzol reagent (Life Technologies). The extracted RNA samples underwent DNase treatment using Turbo DNase (Life Technologies), and RNA-sequencing libraries were prepared using the Illumina stranded total RNA Prep, Ligation with RiboZero Plus kit (Illumina) following the manufacturer’s instructions. The pooled libraries were sequenced in one P3 flow cell on an Illumina NextSeq2000 sequencer using a 100 bp paired-end read protocol. Reads were aligned using STAR/2.7.10a to the GRCh38, GRCm38, and HetGla female 1.0 human, mouse, and naked mole-rat references, respectively.

### Laser-capture microdissection and whole-genome sequencing

Fresh-frozen, OCT-embedded naked mole-rat and mouse dorsal skin samples were cryosectioned at 8 µm thickness and mounted onto Zeiss or Leica PEN-membrane glass slides. Sections were fixed and dehydrated in chilled 70% ethanol, followed by graded ethanol at room temperature, and air-dried for 5 min. Slides were stored at −80 °C in sealed tubes until use. For staining, sections were incubated in hematoxylin (Sigma) for 4 min, rinsed in distilled water, and counterstained with eosin Y (Sigma) for 37 s. Eosin Y solution was prepared by 10-fold dilution in Tris–acetic acid buffer (0.45 M, pH 6.0). Sections were washed twice in 70% ethanol and once in 100% ethanol (10 s each), then air-dried at room temperature for 5 min. All staining procedures were performed at room temperature.

To obtain a comparable number of ∼80 basal epidermal cells across species and treatments (skins of equal thickness), ∼800 µm of interfollicular epidermis along the basement membrane was micro-dissected using two to three micro-biopsies (20X objective, LMD7, Leica). In mice, interfollicular epidermis was across hair follicles. Pooled micro-biopsies were visually inspected in the collection tube lid prior to storage at −80 °C until DNA isolation.

Low-input whole-genome sequencing libraries were generated from LCMD samples following Ellis *et al.* (2021) ^48^. Briefly, LCMD skin samples were digested with Arcturus PicoPure Proteinase K (Thermo Fisher), and genomic DNA was purified using AMPure XP beads (Beckman Coulter). Owing to the limited DNA yield, library preparation was performed immediately using the NEBNext Ultra II FS DNA Library Prep Kit for Illumina (New England Biolabs) in low-input mode (<100 ng), according to the manufacturer’s instructions.

Matched whole-genome sequencing libraries were generated from liver DNA. A small amount of DNA was isolated following the WES protocol, and WGS libraries were prepared using the >100 ng input mode of the NEBNext Ultra II FS DNA Library Prep Kit, consistent with the skin sample workflow. Libraries were pooled and sequenced on an Illumina NovaSeq X 25B platform in paired-end 100 bp mode.

### Single-cell cell-type annotation

The droplet-based sequencing data were aligned using the Cell Ranger Single-Cell Software Suite (v. 3.0.1, 10x Genomics) against the GRCm38 (mm10) and HetGla female 1.0 mouse and naked mole-rat references, respectively. Cells with fewer than 600 detected genes, greater than 10% mitochondrial reads, or log10 total mRNA counts greater than two standard deviations from average across all cells per biopsy were removed.

In the single-species analyses, expression was first normalised using SCtransform ^67^, regressing out the percentage of mitochondrial reads within each biopsy. The different biopsies were then integrated using the canonical correlation analysis (CCA) and mutual nearest neighbour (MNN) approach implemented in the Seurat R package ^68^, using the 3000 most variable features. Doublets were identified separately in each species using the function doubletCells from the scran package ^69^. Cells with a log10 (score + 1) greater than 4 were removed. Cell cycle phase was assigned via the CellCycleScoring Seurat function. Louvain clusters were determined with resolution parameter 0.5 and cluster markers were determined with the FindAllMarkers Seurat function using default parameters.

For the 3-species integration analyses, the count matrices were first subset to one-to-one-to-one orthologue genes. Individual biopsies from all three species were integrated together using the Seurat reciprocal principal component analysis (rPCA) approach using the biopsies with the largest number of cells from each species and each treatment as the reference set. Clusters and markers were determined as above.

The annotation of the high-level cell-types (lymphoid-derived immune cells, myeloid-derived immune-cells, epithelial, fibroblasts, endothelial, muscle, melanocytes, mast, and Schwann cells) was performed in four steps: first, we manually assigned cell-types based on the cluster markers separately for each species (the single-species approach described above). Second, we integrated the three species and manually assigned cell-types based on the cluster markers (the integrated approach described above). Third, for a minority (5.4%) of naked mole-rat cells, the single-species and integrated annotations yielded different results. To break the tie for these ambiguous cells, we generated a third annotation using Garnett with the “3-species-integrated” cell type level 1 marker genes listed in Extended Data Table 3. Finally, we used a 2-layer fully connected feedforward neural network trained on the cells with agreeing single-species and integrated annotations to generate a fourth annotation for the ambiguous cells. For each cell, the final annotation was picked as the consensus of all four methods. Cells for which there was no consensus (2.9 % of all cells) were excluded.

For the sub-clustering of the epithelial compartments, we isolated epithelial cells and re-ran the Louvain clustering with resolution parameter 0.5. For the sub-clustering of fibroblasts, we isolated these cells and re-ran the Louvain clustering with resolution parameters 0.25, 0.5 and 1. For the sub-clustering of myeloid-derived immune cells, we isolated these cells and re-ran the Louvain clustering with resolution parameter 2. Because downstream analyses are potentially sensitive to parameter selection, cluster identities for cell sub-types were manually assigned using multiple marker genes and were numbered rather than collapsed in cases where more granular labelling was not possible (Extended Data Table 3; cell type level 2 markers).

To evaluate the robustness of our annotated cell identities, we conducted a standardized bootstrap-based clustering stability analysis across epithelial, fibroblast, and myeloid-derived immune cell compartments in both mouse and naked mole-rat. The data for each cell type and species were normalized, variable features were selected, the matrix was scaled, and 30 principal components (PCs) were computed. Using these PCs, a nearest-neighbour graph was constructed with k = 20, and Louvain clustering was performed at a resolution of 0.5. Cell type annotations as detailed above served as the fixed baseline against which the stability of re-clustering could be quantified. Robustness was assessed through repeated bootstrap subsampling. For each parameter tested, 50 independent subsampling rounds were performed, each retaining 80% of cells. Agreement between the original and re-clustered labels was quantified using the Jaccard index. Three independent parameter sweeps were carried out: resolution values of 0.2, 0.5, 0.8, and 1.0; PC values 10, 20, and 30; and k-parameter values of 10, 20, 30, and 40.

For the clustering of myeloid-derived immune cells, we isolated these cells and ran the Self-Assembling Manifolds mapping algorithm on their full transcriptomes, correcting for batch effect ^70,71^. We then applied Leiden clustering (resolution = 2) to the resulting cross-species embedding and annotated cluster identities by comparing expression profiles to myeloid immune cell subtype-signatures assembled from publicly available datasets (Extended Data Table 3) ^72^. Due to the limited availability of 1:1 orthologous lymphoid-derived immune cell markers between mouse and naked mole-rat, we independently re-clustered the lymphoid immune cell populations for each species using the *samalg* module of SAMap. Lymphoid immune cell subtypes were annotated manually based on multiple marker genes (Extended Data Table 3).

### Single-cell trajectory analysis of interfollicular epithelial cells

An epithelial cell differentiation trajectory was calculated using Monocle3 on the Seurat-generated UMAP. Its starting point was selected based on the interfollicular epithelial cells containing the highest expression of the undifferentiated basal marker genes KRT14, COL17A1, ITGA6, THY1.

### Differential expression

To perform differential expression (DE) analysis from single-cell data, we summed counts over all cells within each biopsy and subtype to create pseudo-bulk expression profiles. The pseudo-bulk expression profiles of the three species were normalised together (using 1:1:1 orthologue genes), and pairwise differential expression was performed using the DESeq2 package ^73^, with default parameters. For DE genes across epithelial states in Fig. 3 and Extended Data Fig. 4 we created pseudobulks of all IFE cells, Cycling, Undifferentiated state I, II and III, Hybrid state I and II, Differentiated state I-VII. Genes significant in both species (adjusted *P* < 0.05) were retained. Gene expression values were capped at ±4 for visualization. Heatmaps were generated using the pheatmap package [Kolde R (2025). pheatmap: Pretty Heatmaps. R package version 1.0.13; https://cran.r-project.org/package=pheatmap]. P values were adjusted for multiple testing using Benjamini-Hochberg’s method.

Similarly, DE analysis of bulk RNA-sequencing samples was performed using DESeq2 with default parameters. For comparisons across species, only 1:1 orthologue genes were used for differential expression and downstream gene set analysis.

To compare log_2_ fold change differences (Extended Data Fig. 11p), we calculated a linear model using mouse co-culture as the explanatory variable for naked mole-rat co-culture effect (slope [m] = 0.24).

### Gene set enrichment and cell communication scoring

For gene set enrichment analyses, all genes were filtered by adjusted < 0.05 for differential expression in each comparison and ranked by log_2_ fold change. We then used the weighted approach implemented in GSEA ^74^ via the R package fgsea to test for enrichment of gene sets using 1:1:1 orthologous genes in the MSigD/B hallmark and gene ontology (GO) ^75^, Reactome Pathway ^76^, and KEGG databases ^77^. AUCell ^78^ was used to calculate pathway activity scores within individual cells using 1:1:1 orthologous genes for the following gene sets: Collagen biosynthetic process (GO:0032964), Activation of immune response (GO:0002253), and Regulation of T cell activation (GO:0050863), Necroptotic process (GO:0070266), Cell death (GO:0008219), and Cellular senescence (GO:0090398). Likewise, AUCell scores were computed for the hallmark E2F targets (M5925), for the p53 signaling KEGG pathway (hsap04115), and for the set of genes composing the “Hg-cell-death” signature from Kawamura et al. 2023 ^79^ (Extended Data Table 4).

To assign pathway activity weights (Extended Data Fig. 11q), lists of genes and their contribution weights corresponding to pathways of interest were imported from PROGENy ^80,81^. We then applied the R package decoupleR ^82^ to the bulk RNA-seq SCC25 samples and plotted the pathway activities for each sample scaled within each pathway.

Communication scores between fibroblasts and SCC25 cells were defined as the product of ligand and receptor expression for publicly-available ligand-receptor pairs ^83,84^.

### Cell type and state proportions

To test for changes in cell states and types upon treatment, we calculated the proportion of each cell type in each biopsy and applied a centred log ratio (CLR) transformation. This was used as input for linear mixed-effects models (LMM), implemented via the lme4 R package ^85^. To account for the non-independence of biopsies from the same individual, individual identity was included as random effect where appropriate. P values were calculated from the t statistics associated with the slope of each cell type.

### Principal Component Analysis of single-cell pseudobulks

We generated IFE pseudobulk profiles (Fig. 6m; Extended Data Fig. 10i) from naked mole-rats biopsies from the TPA-DMBA-TPA experiment. Raw counts were normalized to counts-per-million (CPM) and subsequently log1p-transformed. The top 1000 most variable genes were then retained for Principal Component Analysis (PCA). Minimum-volume enclosing ellipses were fitted in the PCA space using the Khachiyan optimization algorithm (parameter expand = 0.05), as implemented in the ggforce R package [Pedersen, T. L. ggforce: Accelerating ‘ggplot2’ (R package version 0.5.0.9000, 2025). https://cran.r-project.org/package=ggforce].

### Whole-genome and whole-exome read processing and alignment

For naked mole-rat paired-end FASTQ files were processed per sample, removing adapter contamination and low-quality bases using Trim Galore [Krueger, F. Trim Galore!, Babraham Bioinformatics, 2021). https://github.com/FelixKrueger/TrimGalore] with default settings for paired-end data. Trimmed reads were aligned to the repeat-masked hetGlaV3 reference genome by employing BWA-MEM ^86^. During alignment, read group information (ID, SM, PL=Illumina) was added via the BWA-MEM -R flag, ensuring compatibility with somatic variant calling. Samples sequenced across multiple lanes were merged from lane-level BAM files into a single coordinate-sorted file with samtools merge ^87^. PCR and optical duplicates were then identified and marked using GATK MarkDuplicates ^88^ with simultaneous collection of duplicate metrics for quality control. For WES samples, additionally hybrid-selection capture metrics were calculated by GATK CollectHsMetrics. For mouse paired end FASTQ files were processed by the DKFZ One Touch Pipeline (OTP) with the AlignmentAndQCWorkflow [DKFZ ODCF. AlignmentAndQCWorkflows (2025). https://github.com/DKFZ-ODCF/AlignmentAndQCWorkflows] employing the Roddy workflow management system [TheRoddyWMS. Roddy (2025). https://github.com/TheRoddyWMS/Roddy].

### Somatic variant calling

Somatic variants were called using GATK Mutect2. For skin samples, variant calling was performed in tumor-normal mode, with liver samples from matched animals serving as germline controls, while fibroblast samples were analysed in tumor-only mode. Per-sample VCF files as well as Mutect2 statistics files, generated per chromosome were merged into single per-sample files using GATK MergeVcfs and MergeMutectStats. Read-orientation artefact priors were learned using GATK LearnReadOrientationModel based on the per-chromosome F1R2 archives. The merged somatic VCFs were then processed with GATK FilterMutectCalls to assign FILTER labels, incorporating the merged statistics and the learned read-orientation artefact priors (--ob-priors).

Variants were further annotated via the Ensembl Variant Effect Predictor (VEP) ^89^ to assign predicted functional consequences and affected genes. Annotation was performed in VCF mode using the reference genome FASTA and a matching gene annotation (GTF). Mouse genes were annotated using Ensembl release 102 (mm10), and naked mole-rat genes using mHetGlaV3 (Zenodo v1.4).

For all downstream analyses, only autosomal single-nucleotide variants (SNVs) passing all MuTect2 filters (FILTER = PASS) with a minimum sequencing depth of 5 reads were retained. Whole-exome sequencing (WES) calls were additionally restricted to annotated exonic regions. Variants were further filtered to include only those with a variant allele fraction (VAF; gt_AF) between 0.02 and 0.65.

### Mutation burden analysis

To calculate mutation burden (SNVs per Mb), the number of callable bases was determined per-sample using samtools depth (-q 10 -Q 20), applying standard mapping and base quality thresholds. Callable bases were restricted to autosomal regions and for WES data, to exonic regions and required a minimum sequencing depth of 5 reads in the tumor sample as well as the matched normal. Coverage was additionally capped at the 99th percentile to exclude excessively high-depth regions. To quantify mutation burden in non-synonymous variants synonymous variants were excluded, retaining only variants with predicted harmful functional impact classified as MODERATE or HIGH. To assess mutation patterns in cancer-related genes, we defined a gene list as the union of known skin cancer-associated genes ^56^ and Cancer Gene Census genes ^90^ whose annotated tumor types included the following skin-related keywords: “melanoma,” “SCC,” “basal cell,” “skin cancer,” “cutaneous,” “Spitzoid tumor,” “skin squamous cell,” “skin SCC,” “desmoplastic melanoma,” “skin basal cell carcinoma,” “skin squamous cell carcinoma,” “melanocytic nevus,” and “skin.

### Stem and progenitor replacement rate

Assuming neutral competition, the probability *P* of finding a clone of size *n* progenitors is given by the following equation ^66^:

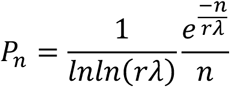

where *r*λ is the loss/replacement rate of the basal progenitor cells. This equation can be manipulated by calculating the first incomplete moment μ of the clone size distribution, defined as:

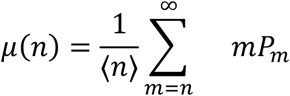

where 〈*n*〉 is the average clone size in a biopsy. If the dynamics of cell loss and replacement follow neutral assumptions, the first incomplete moment assumes an exponential distribution:

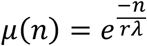

with the corresponding distribution of clone areas A given by

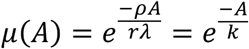

The clone areas are proportional to the biopsy size (1 mm^2^) and the variant allele frequency (VAF) associated with the individual single nucleotide variant that defines the clone. Very few mutations involve a change in copy number, so the areal contribution of individual mutant clones can be approximated as twice the VAF.

The first incomplete moment of cell populations under drift will follow an exponential distribution, so plotting the logarithm of the incomplete moment versus clone area will result in straight lines. For biopsies in which this was observed, the progenitor replacement rate was estimated by fitting the exponent 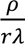 to the incomplete moment distribution.

### Statistical analyses and reproducibility

Statistical tests used are indicated in the figure legends of the corresponding plots.

## Graphical illustrations

Mouse and naked mole-rat illustrations were created by Mikaela Behm and Angela Goncalves.

## Data availability

Mouse and naked mole-rat single-cell RNA-seq datasets (10X Chromium) are available from the European Nucleotide Archive under the accession numbers E-MTAB-12451, E-MTAB-12454, E-MTAB-12456, E-MTAB-12459, E-MTAB-12461, E-MTAB-13556, E-MTAB-16496, E-MTAB-16497, and E-MTAB-16499. The human single-cell RNA-seq dataset is available from the Gene Expression Omnibus (GEO) under the accession number GSE130973. Bulk RNA-seq datasets of co-cultured naked mole-rat and mouse fibroblasts with SCC25 cells are available from the European Nucleotide Archive under the accession number E-MTAB-13538. Mouse and naked mole-rat whole-genome sequencing (WGS) and whole-exome sequencing (WES) datasets are available from the NCBI Sequence Read Archive under the accession number PRJNA903661.

## Acknowledgments

We especially thank Hayley Forest, Alexandra Robinson, and Gemini Chu (Combined Animal Facility, University of Cambridge), as well as Michael Webb and Sarah Elderkin (Cancer Research UK Cambridge Institute), for invaluable support with naked mole-rat skin carcinogenesis experiments. We also thank Greta Döking and colleagues at the University of Duisburg-Essen for their support with naked mole-rat experiments.

We acknowledge the outstanding support and expertise of the DKFZ core facilities, including the NGS, Light Microscopy, Animal, Carcinogen, and Flow Cytometry Core Facilities, as well as the Single-Cell Open Lab. We especially thank Damir Krunic and colleagues at the Light Microscopy Facility, Karin Müller-Decker for assistance with animal experiments, and Frank Lyko for providing single-cell sequencing data.

We thank the Center for Model System and Comparative Pathology, Institute of Pathology, University Hospital Heidelberg, for assistance with tissue processing and DNA damage staining.

We thank Raphael Teodoro Franciscani Coimbra for support with genomic data deposition and data management. We also thank Kimi Pelissero for assistance with DNA extraction from tissue samples.

We thank Ralf Küppers, Jürgen Becker and colleagues at the University of Duisburg-Essen, Faculty of Biology, for support with 10x Chromium experiments, and Radim Šumbera (University of South Bohemia in České Budějovice) for providing juvenile naked mole-rats. We thank Vladimir Botchkarev and Andrei Mardaryev for providing naked mole-rat tissue for training purposes; no data from these tissues were used in this study.

This work was funded by the Helmholtz Association. M.B. was supported by an International Postdoctoral Fellowship from the Swedish Research Council (2017-06173) and a Marie Skłodowska-Curie Individual Fellowship (896324). Additional support was provided by NCT/Helmholtz core funding (A350 to M.F., B270 to D.T.O., B210 to A.G.), the European Research Council (788937 to D.T.O.), a Wellcome Investigator Award (202878/Z/16/Z to D.T.O.), a Helmholtz Junior Group Leader Award (to A.G.), and the Dunhill Medical Trust (RPGF2002/188 to E.St.J.S.).

## Author contributions

MF, AG, DTO, MB: designed experiments.

MF, AG, DTO, MB, VFB: wrote manuscript.

MB, SD: performed experiments.

MB, SW, PZ, PBC, VFB, MF, LPD, AG, NH, JBB: processed and analysed data.

MB, LP, MK, SDP, SBe, MBe, BB, DS, MS, CR, JD, FC, REW: collected and processed tissue samples.

FL: curated data and provided data management.

SBl, REW: provided mouse imaging data.

SBe, ESJ, WTK: provided naked mole-rat tissues or cells.

LSB: provided human skin single cell RNA sequencing data.

ASL: provided human skin samples.

## Declaration of interests

All authors declare no conflict of interest.

---

**Extended Data Fig. 1:**
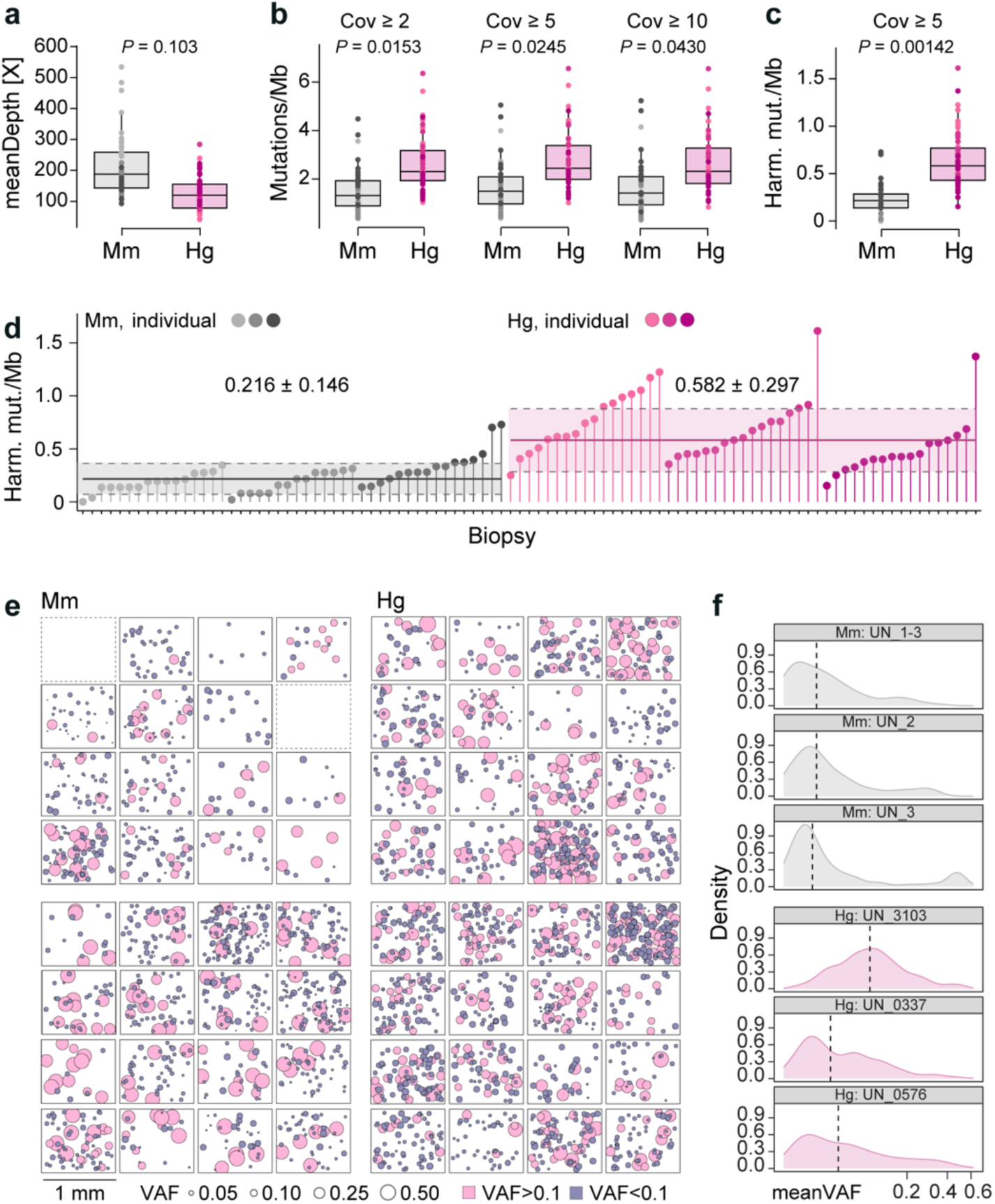
Somatic mutations and clonal dynamics in naked mole-rat and mouse skin. **a**, Mean whole-exome sequencing depth of skin biopsies from mouse (*Mm*) and naked mole-rat (*Hg*). **b**, SNV mutation burden across coverage thresholds. **c**, **d**, Harmful somatic SNV burden at coverage ≥5 (**c**) and per biopsy (lollipop) (**d**). Biopsies are grouped by individual. Solid line indicates the median; dotted line indicates ± s.d. **e**, Nonsynonymous SNVs with predicted moderate or high functional impact in *Mm* (left) and *Hg* (right) biopsies. Each square represents one biopsy; each circle represents one mutated gene; circle size indicates VAF (blue, <0.1; pink, >0.1). Dotted squares indicate no mutated gene passing filtering criteria. **f**, Distribution of harmful nonsynonymous SNVs per individual shown on a pseudo-logarithmic scale. Dashed lines indicate median VAF. Box plots show the median and interquartile range, with 1.5× IQR whiskers (**a–c**). *Mm*, N = 3, n = 14-16; *Hg*, N = 3, n = 17 (**a–d**, **f**). *Mm*, N = 2, n = 14-16; *Hg*, N = 2, n = 16 (**e**). Linear mixed-effects model with individual as a random effect; two-sided likelihood-ratio tests (**a–c**).

**Extended Data Fig. 2:**
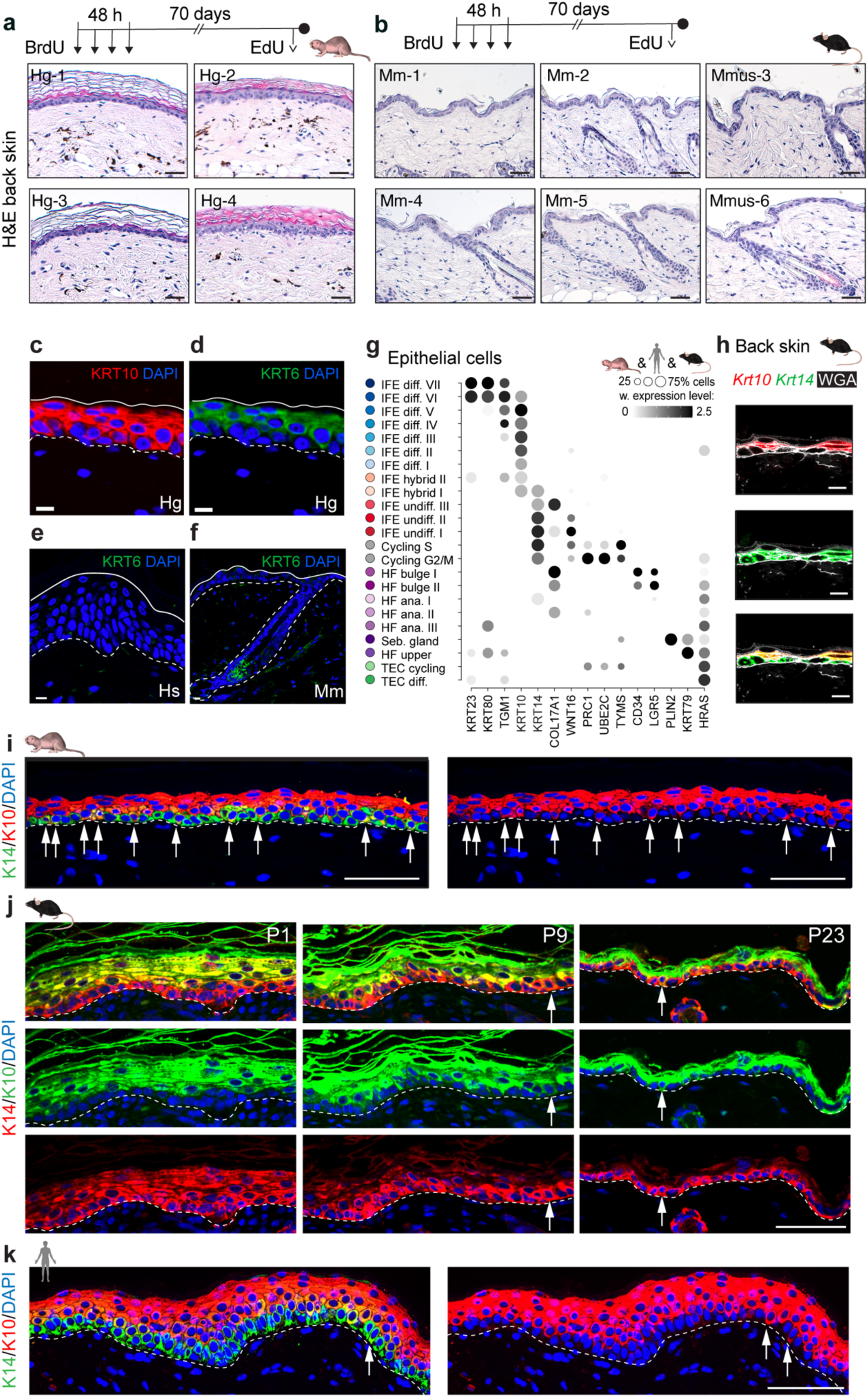
Long-term labelling of epidermal cells and identification of hybrid epidermal cells in naked mole-rat skin. **a**, **b**, H&E staining of naked mole-rat (*Hg*) (**a**) and mouse (*Mm*) (**b**) back skin 70 days after four BrdU injections. **c–f**, KRT6 immunofluorescence in *Hg* (**c d**), human (*Hs*, **e**) and *Mm* (**f**) back skin. **g**, Dot plot of orthologous gene expression used to classify epithelial cell types in *Hg*, *Hs* and *Mm* skin. Colour intensity indicates expression level; dot size indicates the percentage of expressing cells. IFE, interfollicular epidermis; diff., differentiated; undiff., undifferentiated; HF, hair follicle; ana., anagen; Seb., sebaceous; TEC, tumour epithelial cell. **h**, RNAscope labelling of *Krt10* and *Krt14* in *Mm* back skin. **i–k**, Immunofluorescence of KRT14 and KRT10 in *Hg* (**i**), *Mm* (**j**) and *Hs* (**k**) skin. WGA labels the plasma membrane (**h**). Dotted lines indicate the basement membrane (**c–f**, **i–k**); solid lines indicate the outer cornified layer (**c–f**). P1, P9 and P23 denote postnatal days 1, 9 and 23 (**j**). Arrows indicate basal KRT14⁺KRT10⁺ cells (**i–k**). Scale bars, 50 μm (**a**, **b**), 10 μm (**c–f**, **h**) and 100 μm (**i–k**).

**Extended Data Fig. 3:**
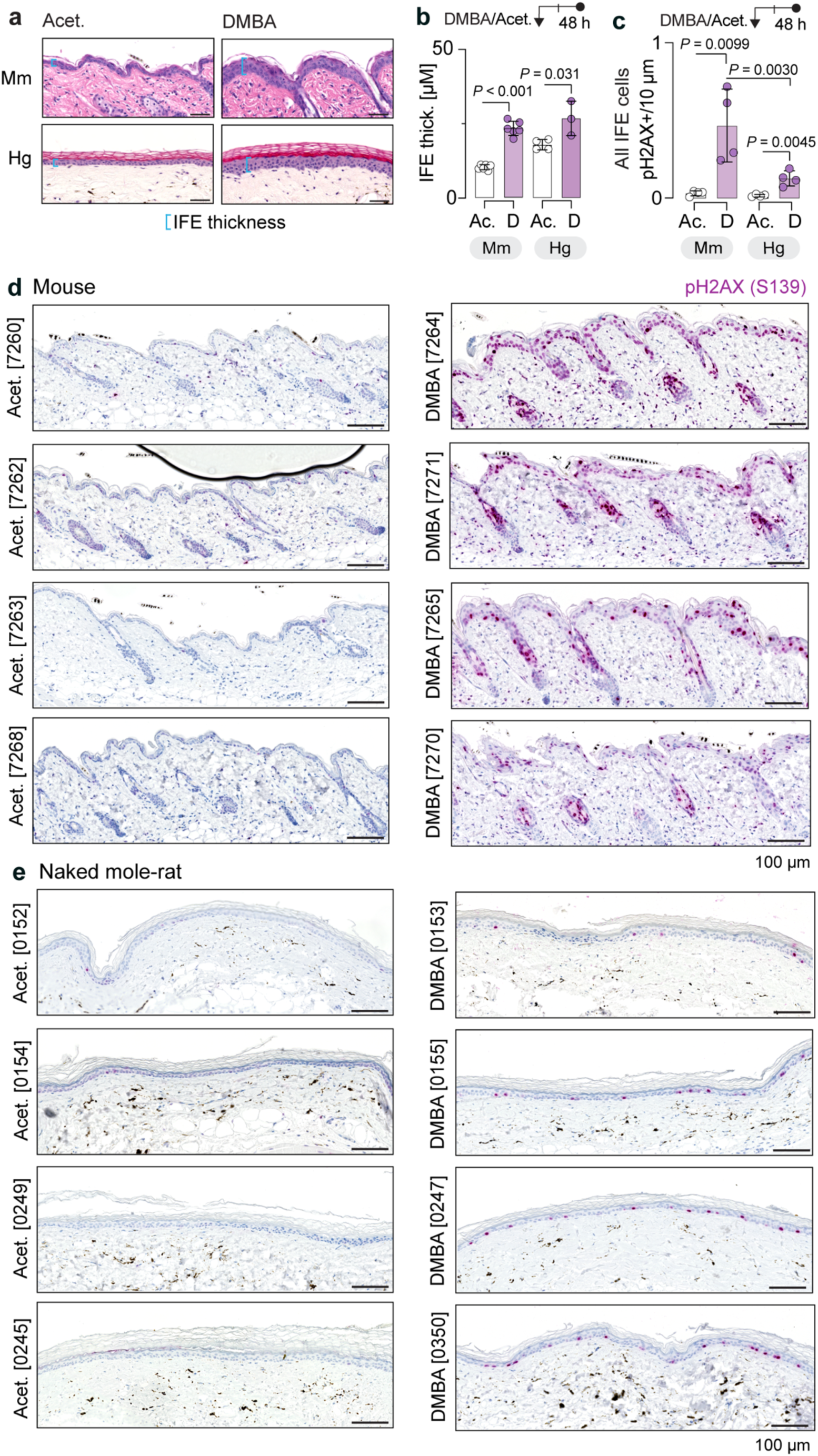
DMBA-induced DNA damage response in mouse and naked mole-rat skin. **a**, **b**, H&E-stained sections of mouse (*Mm*) and naked mole-rat (*Hg*) back skin 48 h after topical DMBA (D) or acetone vehicle (Acet., Ac.) (**a**) and quantification of IFE thickness (**b**). **c–e**, Quantification of pH2AX (S139)^+^ IFE cells in *Mm* and *Hg* back skin 48 h after DMBA or acetone treatment (**c**), with representative pH2AX-stained sections shown in **d**, **e**. Scale bars, 50 μm (**a**) and 100 μm (**d**, **e**). Data points represent individual means **(b**, **c)**. Values show mean ± s.d. (**b**, **c**). N = 6 biologically independent individuals for *Mm* (Ac., and D), N = 4, for *Hg* Ac. N = 3 and for D (**b**); N = 4 biologically independent individuals per species and treatment (**c**). Unpaired two-tailed *t*-test (**b**, **c**).

**Extended Data Fig. 4:**
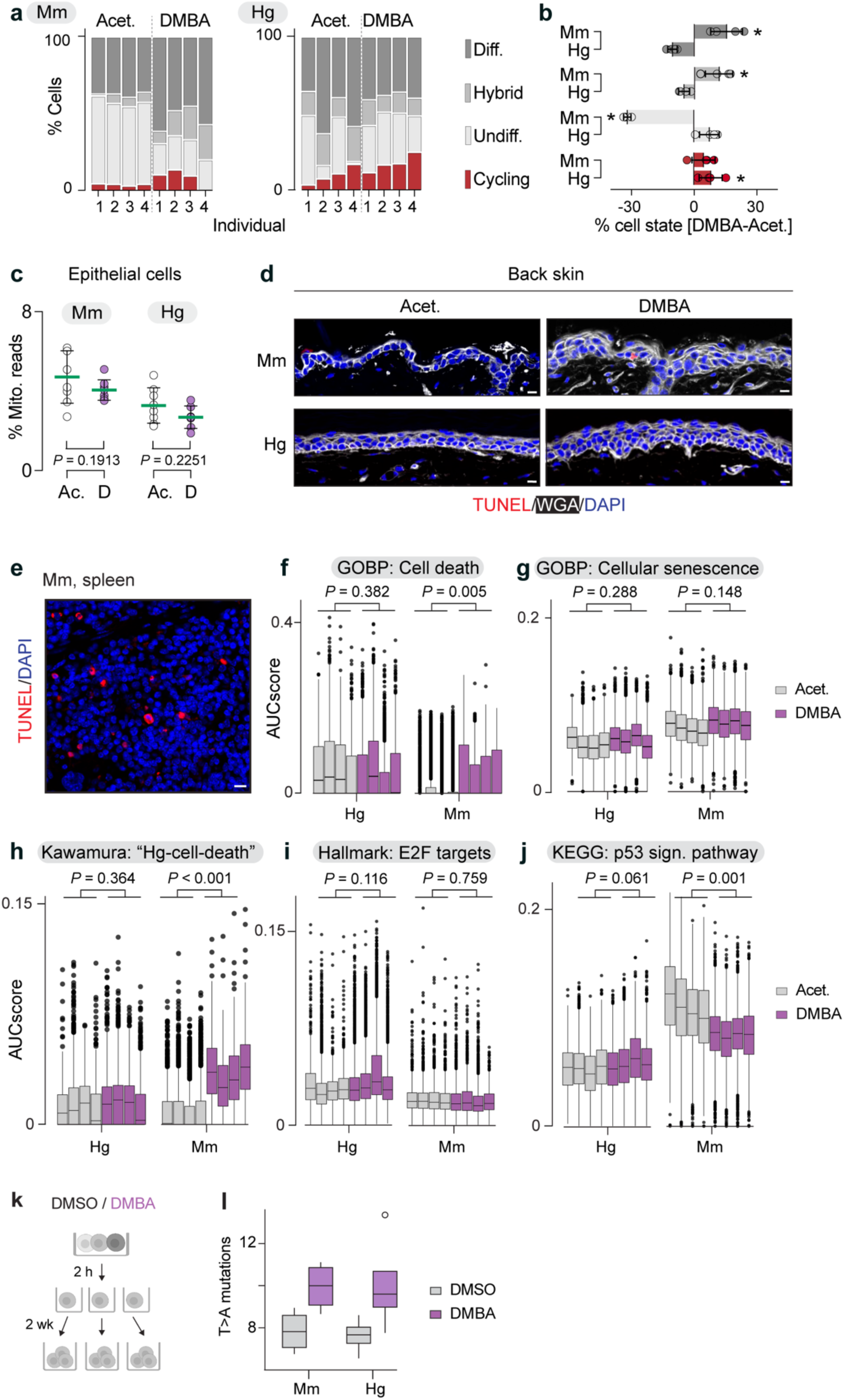
Differentiation and cell death responses to DMBA-induced mutagenic stress. **a**, Percentage of cycling, undifferentiated (Undiff.), hybrid and differentiated (Diff.) IFE cells 48 h after topical acetone (Acet.) or DMBA treatment in mouse (*Mm*, left) and naked mole-rat (*Hg*, right) skin. **b**, DMBA-induced change in IFE cell-state proportions in *Mm* and *Hg* skin relative to acetone controls. **c**, % of mitochondrial RNA reads in *Mm* and *Hg* IFE cells 48 h after acetone (Ac.) or DMBA (D) treatment. **d**, **e**, TUNEL staining of *Mm* and *Hg* back skin (**d**) and *Mm* spleen (**e**). **f–j**, Gene set scoring of cell death**–**related gene sets in *Mm* and *Hg* IFE cells. **k**, **l**, Clonal expansion of DMSO vehicle- or DMBA-treated *Mm* and *Hg* fibroblasts *in vitro* (**k**) and percentage of T>A mutations (**l**). Scale bars, 10 μm (**d**, **e**). Values show mean ± s.d. (**b**, **c**). Box plots show median, interquartile range, with 1.5× IQR whiskers (**f–j**). Bars represent one biopsy from one individual (N = 4 per species and treatment) in (**a**), and the mean of these biopsies in (**b**). Data points and boxes represent individual biopsies (**c**, **f–j**). n = 4 independently derived single-cell clones per condition and species (**l**). Linear model (**b**); linear mixed-effects model (**c**, **f–j**). \**P* < 0.05 (**b**). Exact *P* values are shown (**c**, **f–j**).

**Extended Data Fig. 5:**
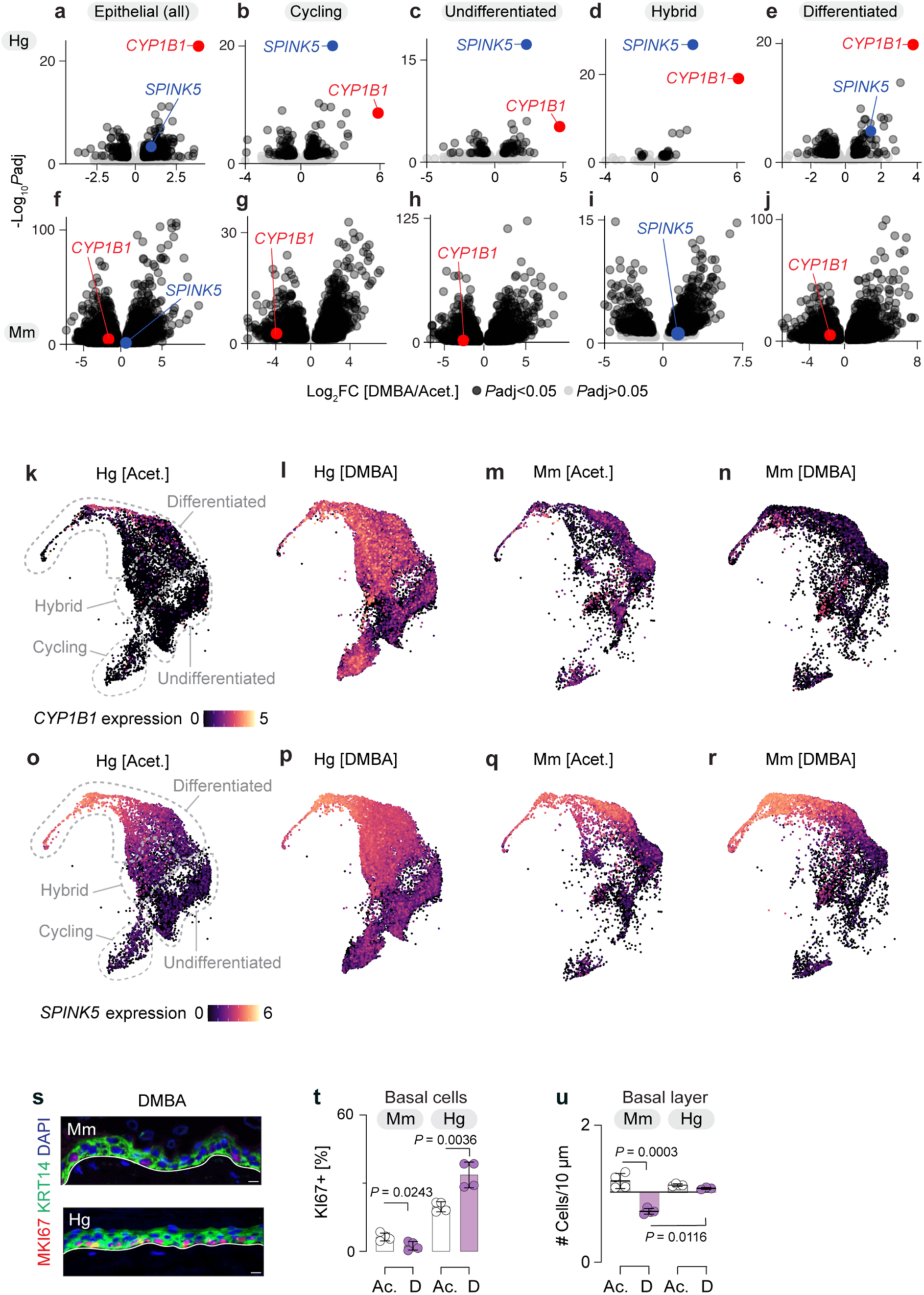
Mutagen-induced transcriptional changes in mouse and naked mole-rat skin. **ya–j**, Volcano plots of differentially expressed (DE) genes from scRNA-seq of naked mole-rat (*Hg*; **a–e**) and mouse (*Mm*; **f–j**) IFE treated with DMBA or acetone (Acet.). DE analysis was performed in all IFE cells (**a**, **f**), cycling cells (**b**, **g**), undifferentiated IFE cells (**c**, **h**), hybrid IFE cells (**d**, **i**), and differentiated IFE cells (**e**, **j**). Genes with *P*adj < 0.05 are shown in black and *P*adj > 0.05 in light grey. *CYP1B1* (red) and *SPINK5* (blue) are highlighted where *P*adj < 0.05. **k–r**, UMAPs showing expression of *CYP1B1* (**k–n**) and *SPINK5* (**o–r**) in DMBA- and acet.-treated *Hg* and *Mm* IFE cells. **s**, **t**, Immunofluorescence images (**s**) and quantification (**t**) of MKI67^+^ cells per 100 basal cells in DMBA (D)- or acetone (Ac.)-treated *Mm* (top) and *Hg* (bottom) skin. Scale bars, 10 μm (**s**). **u**, Number of cells per 10 μm basal epidermal layer in *Mm* and *Hg* Ac.- or D-treated skin (**u**). Values show mean ± s.d. (**t**, **u**). N = 4 biologically independent individuals per species and treatment (**a–r**, **t**); N = 3–4 biologically independent individuals per species and treatment (**u**). Unpaired two-tailed *t*-test (**t**, **u**).

**Extended Data Fig. 6:**
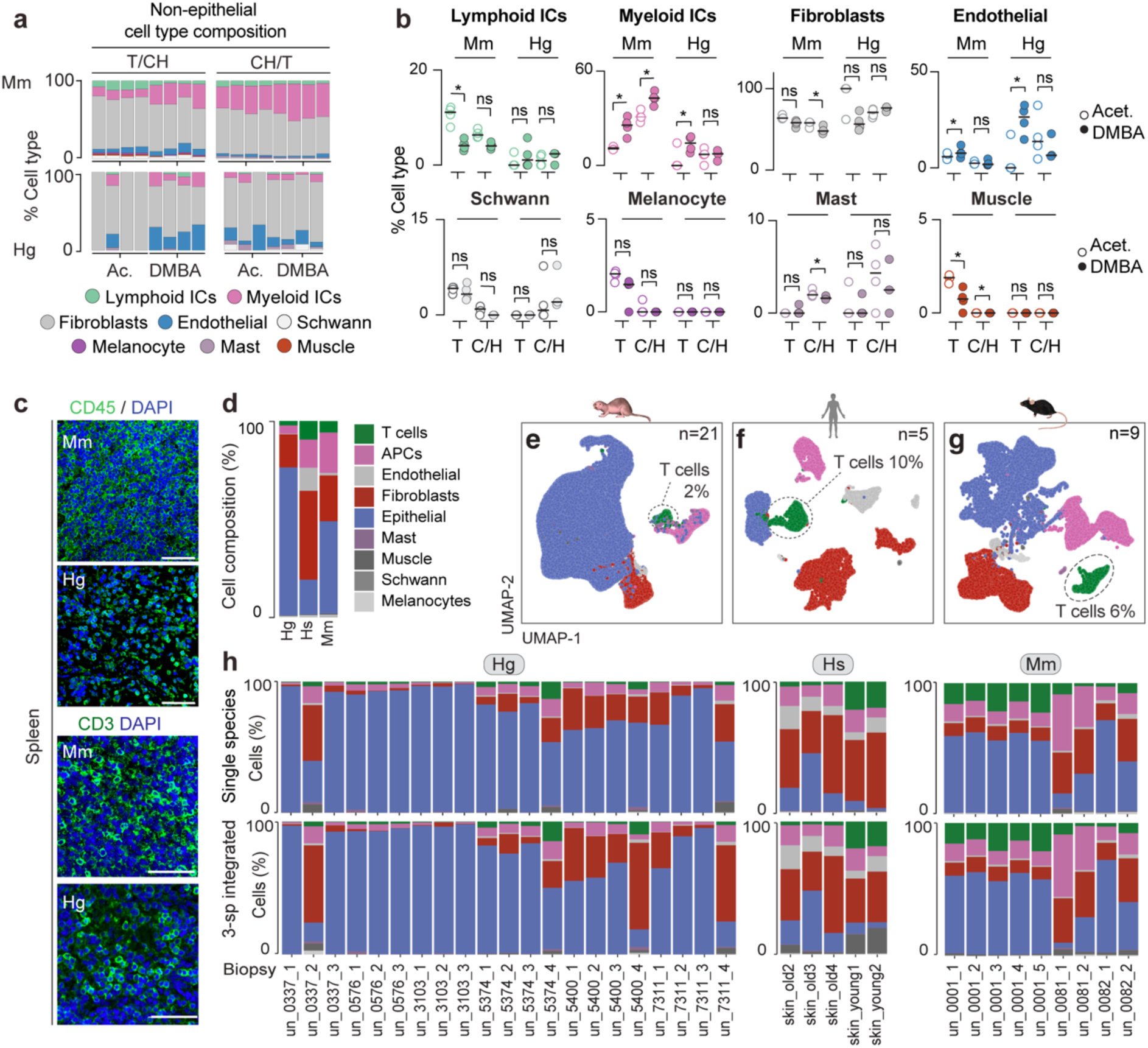
Cross-species characterisation of skin cell composition. **a**, Cell-type composition per biopsy in the non-epithelial skin compartment of mouse (*Mm*, top) and naked mole-rat (*Hg*, bottom) skin treated with acetone (Ac.) or DMBA for 48 h. Skin digestion order: trypsin followed by collagenase–hyaluronidase (T/CH) or collagenase–hyaluronidase followed by trypsin (CH/T). ICs, immune cells. **b**, Relative abundance of cell types in *Mm* and *Hg* skin biopsies treated with acetone (open circles) or DMBA (filled circles) following T/CH or CH/T digestion. **c**, Immunofluorescence of CD45⁺ immune cells (top) and CD3⁺ T cells (bottom) in *Mm* and *Hg* spleen. Scale bars, 50 μm. **d–g**, Cell-type composition (**d**) and UMAPs (**e–g**) of homeostatic *Hg* (**e**), human (*Hs*, **f**) and *Mm* (**g**) skin. **h**, Proportion of cell types per species (top) or after three-species (3-sp) integration (bottom) in *Hg*, *Hs* and *Mm* skin biopsies. N = 3–4 biologically independent individuals per species, treatment and digestion order (**a**, **b**); *Hg* (n = 21 biopsies from 6 individuals), *Hs* (n = 5 biopsies from 5 individuals), *Mm* (n = 9 biopsies from 2 individuals) (**d–h**). Linear model (**b**). \**P* < 0.05; ns, not significant (*P* > 0.05).

**Extended Data Fig. 7:**
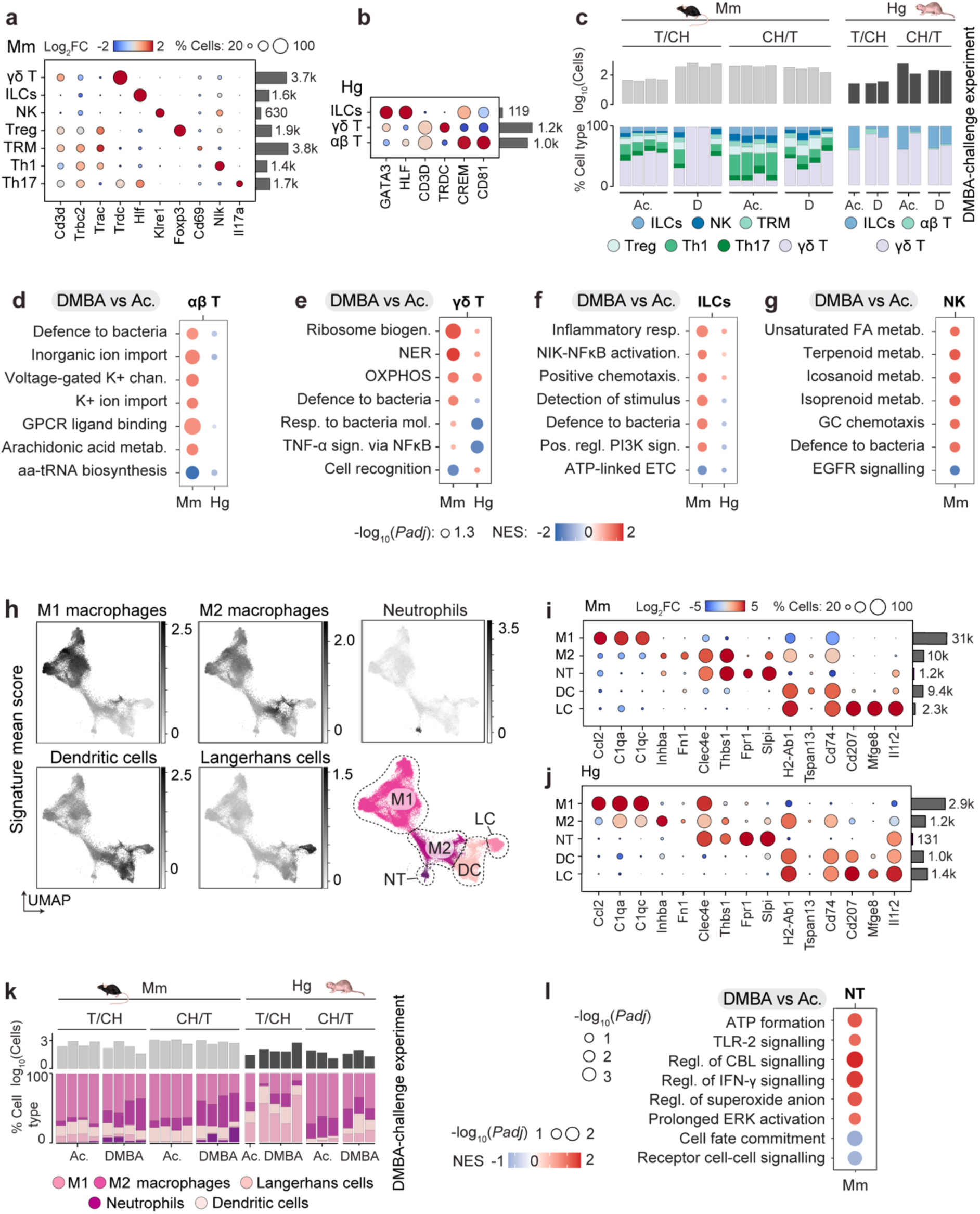
Immune responses to acute carcinogen challenge in mouse and naked mole-rat skin. **a**, **b**, Log₂ fold change of lymphoid immune marker gene expression in mouse (*Mm*) (**a**) and naked mole-rat (*Hg*) (**b**) skin. Dot size indicates the percentage of cells expressing each marker per subtype; numbers indicate cell counts (k, x10³). ILCs, innate lymphoid cells; NK, natural killer cells; Treg, regulatory T; TRM, tissue-resident memory T; Th1/Th17, T helper 1/17. **c**, Log₁₀-transformed counts (top) and relative proportions (bottom) of lymphoid immune cells per biopsy in *Mm* and *Hg* acetone (Ac.)- or DMBA (D)-treated skin for 48 h. Skin digestion order: trypsin followed by collagenase–hyaluronidase (T/CH) or collagenase–hyaluronidase followed by trypsin (CH/T). **d–g**, Gene set enrichment analysis of αβ T cells (**d**), γδ T cells (**e**), ILCs (**f**) and NK cells (**g**) comparing DMBA-versus Ac.-treated *Mm* and *Hg* skin. Circle size reflects significance (*P*adj < 0.05); colour indicates NES. GPCR, G protein–coupled receptor; FA, fatty acid; GC, granulocyte. **h**, Subtype signature scores projected onto UMAP of myeloid immune cells from *Mm* and *Hg* skin. M1/M2, type 1/2 macrophages; DC, dendritic cells; LC, Langerhans cells; NT, neutrophils. **i**, **j**, Log₂ fold change of myeloid immune marker gene expression in *Mm* (**i**) and *Hg* (**j**) skin. Dot size indicates the percentage of cells expressing each marker per subtype; numbers indicate cell counts (k, x10³). **k**, Log₁₀-transformed counts (top) and relative proportions (bottom) of myeloid immune cells per biopsy in *Mm* and *Hg* Ac.- or D-treated skin for 48 h. **l**, Gene set enrichment analysis of NTs comparing DMBA-versus Ac.-treated *Mm* and *Hg* skin. Circle size reflects significance; colour indicates NES. N = 3–4 biologically independent individuals per species and treatment (**a**, **b**, **d–j**, **l**); n = 1–4 biologically independent individuals per species, treatment and digestion order (**c**, **k**).

**Extended Data Fig. 8:**
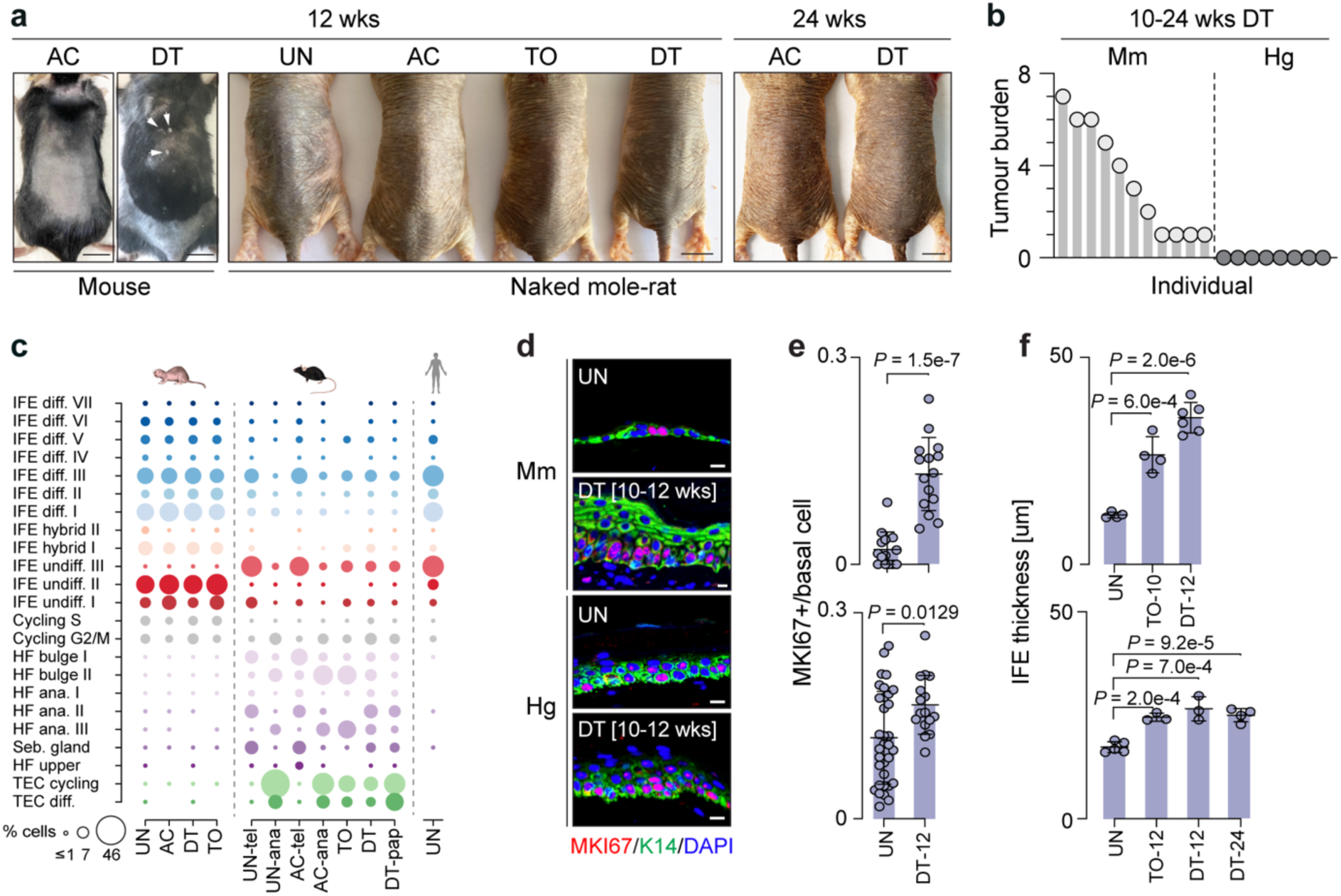
Two-stage chemical carcinogenesis in naked mole-rat and mouse skin. **a**, Images of mice (*Mm*) and naked mole-rats (*Hg*) treated with DMBA/TPA (DT) or respective controls. UN, untreated; AC, acetone; TO, TPA only for 12 or 24 weeks. Arrowheads indicate papillomas. **b**, Tumour burden per DT-treated animal. *Mm*: 10-12 weeks, N = 11; *Hg*: 12 weeks, N = 4; 24 weeks, N = 4. **c**, Mean percentage of cells per IFE differentiation state in *Hg*, *Mm* and human skin. HF, hair follicle; ana., anagen; Seb., sebaceous; TEC, tumour epithelial cell; tel., telogen; pap., papilloma. **d**, **e**, Immunofluorescence (**d**) and quantification (**e**) of MKI67⁺ IFE cells in UN- or DT-treated *Mm* and *Hg* skin (N = 3–4 per condition; data points represent one measurement). **f**, IFE thickness in UN-, TO- (10 or 12 weeks) or DT-treated (12 or 24 weeks) *Mm* and *Hg* back skin. Data points represent one biological replicate averaged from ≥5 images (*Mm*, N = 4–6; *Hg*, N = 3–5). Scale bars, 1 cm (**a**), 10 μm (**d**). Values show mean ± s.d. (**e**, **f**) and mean (**c**). Unpaired two-tailed *t*-test (**e**, **f**). Exact *P* values are shown (**e**, **f**).

**Extended Data Fig. 9:**
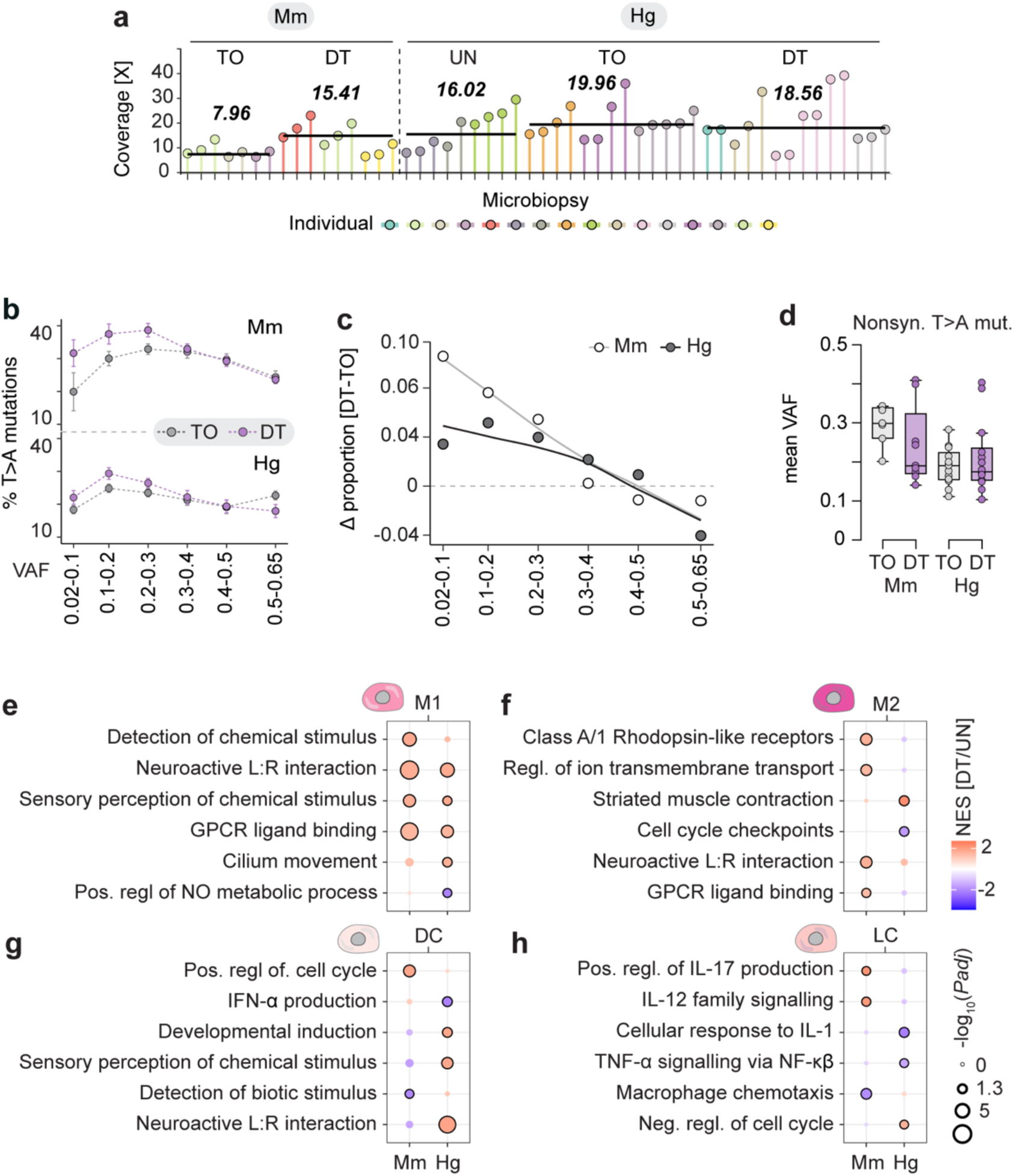
Somatic mutations in carcinogen-treated mouse and naked mole-rat skin. **a**, Whole-genome sequencing coverage of microdissected mouse (*Mm*) and naked mole-rat (*Hg*) IFE regions. Thick line indicates the median per treatment. UN, untreated; TO, TPA only; DT, DMBA/TPA. Each lollipop represents one replicate. **b**, Mean proportion of T>A mutations per VAF bin in TO- and DT-treated *Mm* (top) and *Hg* (bottom) skin. Data points show means across individuals; error bars indicate ± s.e.m. *Mm*: TO, N = 3, n = 2–3; DT, N = 3, n = 3. *Hg*: TO, N = 3, n = 4–5; DT, N = 4, n = 2–6. **c**, Delta T>A SNVs across VAF bins in *Mm* and *Hg* micro-biopsies (DT − TO). Data points show per-bin differences in mean T>A proportion. Sample sizes as in **b**. **d**, Mean VAF of nonsynonymous T>A SNVs in TO- and DT-treated *Mm* and *Hg* skin. Box plots show median and interquartile range, with 1.5× IQR whiskers. Sample sizes as in **b**. **e–h**, Gene set enrichment analysis of M1 macrophages (**e**), M2 macrophages (**f**), dendritic cells (**g**) and Langerhans cells (**h**) comparing DT-versus UN-treated *Mm* and *Hg* skin. Circle size indicates significance; colour indicates NES. L:R, ligand:receptor; GPCR, G protein–coupled receptor; NO, nitric oxide.

**Extended Data Fig. 10:**
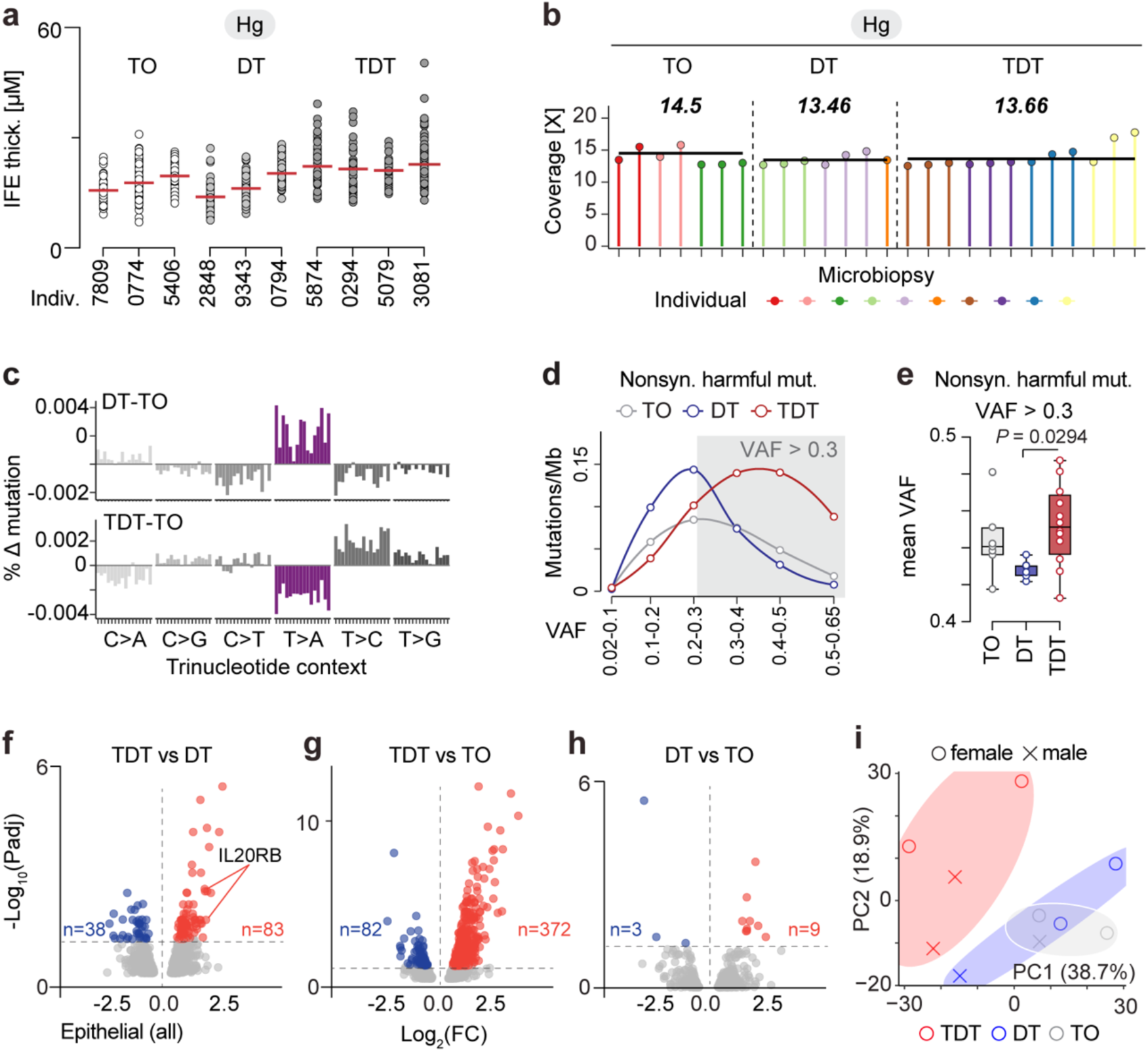
Epidermal priming by TPA enables carcinogen-induced alterations in naked mole-rat skin. **a**, IFE thickness in naked mole-rat (*Hg*) back skin treated with TPA only (TO), DMBA/TPA (DT) or TPA/DMBA/TPA (TDT). Data points represent individual measurements per individual; mean is shown (red line). **b**, Whole-genome sequencing coverage of microdissected IFE regions. Thick line indicates the median per treatment. Each lollipop represents one replicate. **c**, Frequencies of SNVs across 96 trinucleotide contexts in DT- (top) or TDT-treated (bottom) *Hg* IFE regions after subtraction of TO background. **d**, **e**, Nonsynonymous harmful SNV mutation burden by VAF bin (**d**) and mean VAF (**e**). **f–h**, Volcano plots of significantly regulated genes (*P*adj < 0.05) in *Hg* IFE cells comparing TDT versus DT (**f**), TDT versus TO (**g**) and DT versus TO (**h**). **i**, Principal component analysis of pseudo-bulked single-cell transcriptomes from female and male *Hg* IFE cells; shaded areas indicate treatment-specific confidence ellipses. *Hg:* TO, N = 3, n = 2–3; DT, N = 3, n = 1–3; TDT, N = 4, n = 3 (**b–e**). *Hg,* N = 3-4 biologically independent individuals per treatment group (**f–i)**.

**Extended Data Fig. 11:**
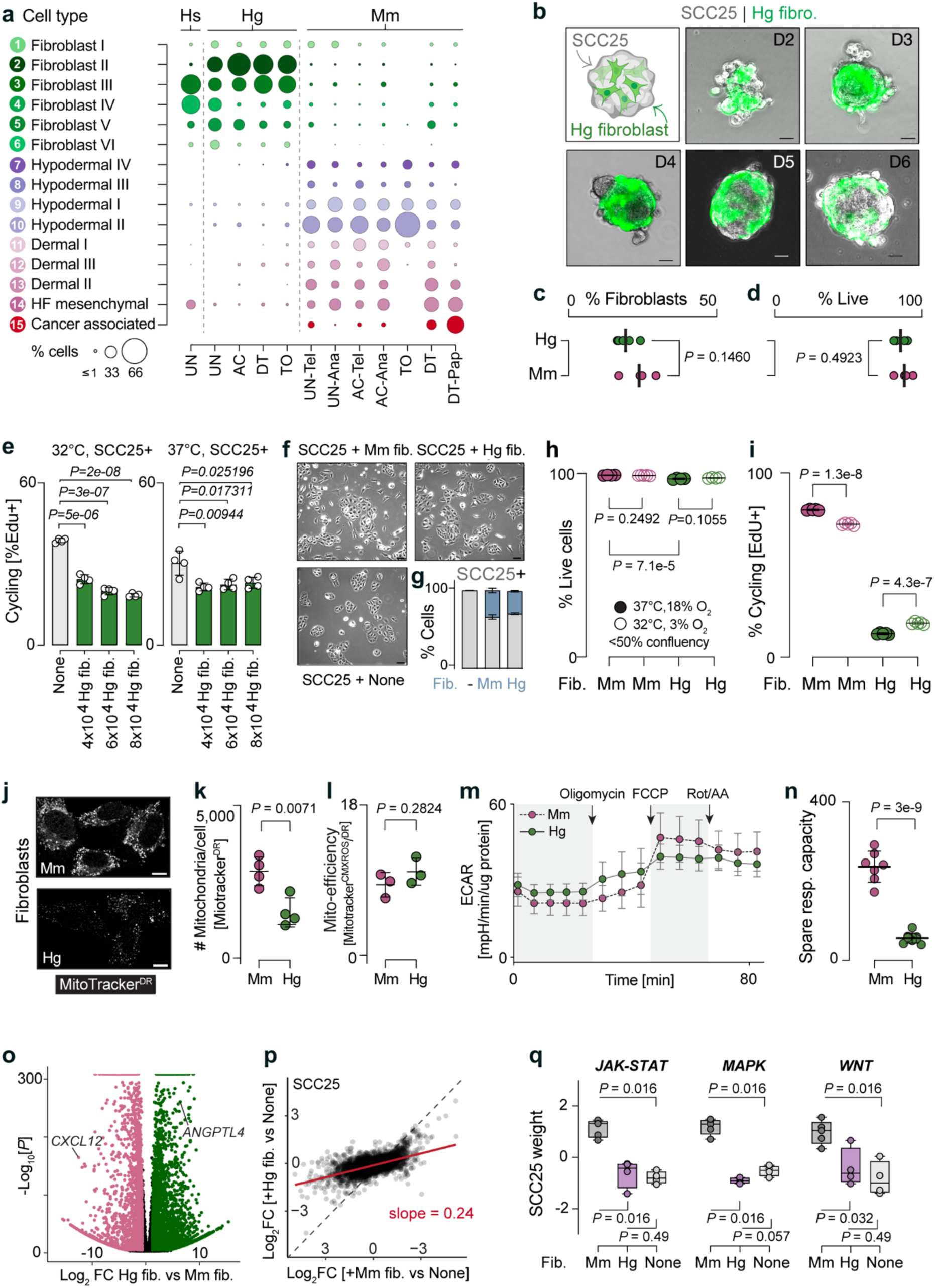
Epithelial growth suppressive features of naked mole-rat fibroblasts. **a**, Skin fibroblasts cell states in naked mole-rat (*Hg*), human (*Hs*) and mouse (*Mm*). UN, untreated; AC, acetone; TO, TPA only; DT, DMBA/TPA; Tel, telogen; Ana, anagen; Pap, papilloma; HF, hair follicle. *Hg*: UN, N = 5, n = 14; AC, N = 3, n = 8; TO, N = 3, n = 8; DT, N = 4, n = 11. *Hs*: N = 5, n = 5. *Mm*: UN–Tel, N = 2, n = 2; UN–Ana, N = 1, n = 1; AC–Tel, N = 3, n = 6; AC–Ana, N = 3, n = 3; TO, N = 3, n = 3; DT, N = 3, n = 5; DT–Pap, N = 2, n = 2. **b–d**, 3D co-culture images (**b**), percentage of fibroblasts (**c**) and viability (**d**) of *Hg* and *Mm* fibroblasts from day (D) 2–6. One representative image from ≥5 images per day is shown (**b**). Data points represent replicates from one representative experiment of two independently performed experiments (**c**, **d**); n = 4–5 per species. **e**, EdU^+^ cycling SCC25 cells in 2D co-culture with increasing numbers of *Hg* fibroblasts at 32 °C (left) or 37 °C (right). Data points represent replicates from one experiment; n = 4 per condition. **f**, **g**, 2D co-culture images (**f**) and cell-type proportions (**g**) after 48 h co-culture of SCC25 alone (None) or with *Mm* or *Hg* fibroblasts. **h**, **i**, Viability (**h**) and proliferation (**i**) of *Mm* or *Hg* fibroblasts cultured at 37 °C and 18% O₂ or 32 °C and 3% O₂. n = 4–5 replicates from one representative experiment of three independent experiments (**f–i**). **j**, MitoTracker Deep Red staining of *Mm* (top) and *Hg* (bottom) fibroblasts. **k–n**, Number of mitochondria per fibroblast (**k**), mitochondrial efficiency (**l**), extracellular acidification rate (**m**) and spare respiratory capacity (**n**) in *Mm* and *Hg* fibroblasts. Data points represent the mean of technical replicates from one representative experiment of three independent experiments (**k**, **l**: n = 3–4; **m**, **n**: n = 8–9). **o**, Volcano plot of differentially expressed orthologous genes in *Hg* versus *Mm* fibroblasts. **p**, Correlation of genes differentially expressed in SCC25 co-cultured with *Hg* (y axis) or *Mm* (x axis) fibroblasts vs SCC25 alone. **q**, Scaled signalling pathway scores in SCC25 cells after co-culture with *Mm*, *Hg* or no fibroblasts. n = 4–5 per species (**o–q**). Scale bars, 10 μm (**b**, **f**, **j**). Values show mean ± s.d. (**c–e**, **h**, **i**, **k–n**). Box plots show quartiles and range (**q**). Unpaired two-tailed *t*-test (**c–e**, **h**, **i**, **k**, **l**, **n**). Wilcoxon test (**q**). Exact *P* values are shown.

**Extended Data Fig. 12:**
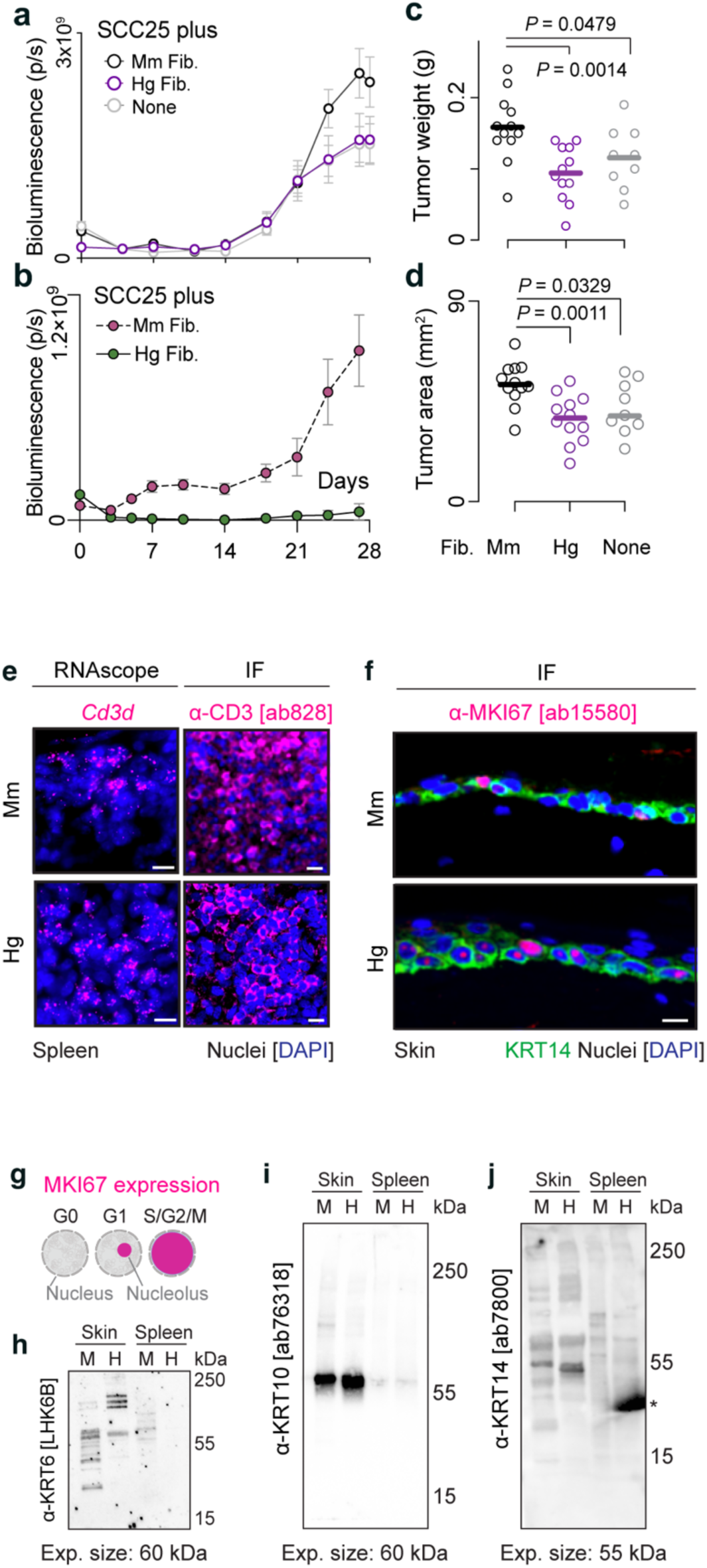
In vivo co-injection assays and antibody validation in naked mole-rat tissues. **a–d**, Quantification of cancer cell (**a**) or fibroblast (**b**) xenograft growth, tumour weight (**c**), and area (**d**) 28 days after co-injection of SCC25 cells with mouse (*Mm*), naked mole-rat (*Hg*), or no fibroblasts (None; Fib.). Data points represent an average of 4–5 mice per condition (**a**, **b**) and measurement from one tumour (**c**, **d**). **e**, RNAscope labelling of *Cd3d* (**e**, left) and CD3 immunofluorescence (IF, **e**, right) in *Mm* and *Hg* spleen. **f**, Immunofluorescence of MKI67 expression in *Mm* and *Hg* skin. **g**, Schematic of MKI67 nuclear localisation and expression across the cell cycle. Cells express MKI67 during the G1, S, G2, and mitotic phases, but not during G0. During early G1, MKI67 localises to the nucleoli. **h–j**, Western blot analysis of KRT6 (**h**), KRT10 (**i**), and KRT14 (**j**) in mouse (M) and naked mole-rat (H) skin and spleen. Exp., expected; *nonspecific binding. Scale bars, 10 μm (**e**, **f**). Values show mean ± s.e.m. (**a**, **b**) and mean ± s.d. (**c**, **d**). Unpaired two-tailed *t*-test (**c**, **d**). Exact *P* values are shown.

## Notes

### Competing Interest Statement

The authors have declared no competing interest.

### Summary of Updates

This version of the manuscript has been revised to incorporate additional experiments and analyses that strengthen the conclusions. The authorship and acknowledgments have been updated.

## References

1 Cagan, A. et al. Somatic mutation rates scale with lifespan across mammals. Nature 604, 517–524 (2022). 10.1038/s41586-022-04618-z

2 Buffenstein, R. et al. The naked truth: a comprehensive clarification and classification of current ‘myths’ in naked mole-rat biology. Biol Rev Camb Philos Soc 97, 115–140 (2022). 10.1111/brv.12791

3 Braude, S. et al. Surprisingly long survival of premature conclusions about naked mole-rat biology. Biol Rev 96, 376–393 (2021). 10.1111/brv.12660

4 Yamamura, Y., Kawamura, Y., Oka, K. & Miura, K. Carcinogenesis resistance in the longest-lived rodent, the naked mole-rat. Cancer Sci 113, 4030–4036 (2022). 10.1111/cas.15570

5 Delaney, M. A., Nagy, L., Kinsel, M. J. & Treuting, P. M. Spontaneous histologic lesions of the adult naked mole rat (Heterocephalus glaber): a retrospective survey of lesions in a zoo population. Vet Pathol 50, 607–621 (2013). 10.1177/0300985812471543

6 Delaney, M. A. et al. Initial Case Reports of Cancer in Naked Mole-rats (Heterocephalus glaber). Veterinary Pathology 53, 691–696 (2016). 10.1177/0300985816630796

7 Taylor, K. R., Milone, N. A. & Rodriguez, C. E. Four Cases of Spontaneous Neoplasia in the Naked Mole-Rat (Heterocephalus glaber), A Putative Cancer-Resistant Species. J Gerontol a-Biol 72, 38–43 (2017). 10.1093/gerona/glw047

8 Cole, J. E., Steeil, J. C., Sarro, S. J., Kerns, K. L. & Cartoceti, A. Chordoma of the sacrum of an adult naked mole-rat. J Vet Diagn Invest 32, 132–135 (2020). Artn1040638719894985 10.1177/1040638719894985

9 Hadi, F. et al. Transformation of naked mole-rat cells. Nature 583, E1-+ (2020). 10.1038/s41586-020-2410-x

10 Zhao, J. et al. Reply to: Transformation of naked mole-rat cells. Nature 583, E8-+ (2020). 10.1038/s41586-020-2411-9

11 Shepard, A. et al. An Autochthonous Model of Lung Cancer Identifies Requirements for Cellular Transformation in the Naked Mole Rat. Cancer Discov 16, 35–45 (2026). 10.1158/2159-8290.CD-25-0526

12 MacRae, S. L. et al. DNA repair in species with extreme lifespan differences. Aging-Us 7, 1171–1184 (2015). DOI 10.18632/aging.100866

13 Kim, E. B. et al. Genome sequencing reveals insights into physiology and longevity of the naked mole rat. Nature 479, 223–227 (2011). 10.1038/nature10533

14 Evdokimov, A. et al. Naked mole rat cells display more efficient excision repair than mouse cells. Aging-Us 10, 1454–1473 (2018). 10.18632/aging.101482

15 Chen, Y. et al. A cGAS-mediated mechanism in naked mole-rats potentiates DNA repair and delays aging. Science 390, eadp5056 (2025). 10.1126/science.adp5056

16 Seluanov, A. et al. Hypersensitivity to contact inhibition provides a clue to cancer resistance of naked mole-rat. Proc Natl Acad Sci U S A 106, 19352–19357 (2009). 10.1073/pnas.0905252106

17 Tian, X. et al. INK4 locus of the tumor-resistant rodent, the naked mole rat, expresses a functional p15/p16 hybrid isoform. Proceedings of the National Academy of Sciences of the United States of America 112, 1053–1058 (2015). 10.1073/pnas.1418203112

18 Tian, X. et al. High-molecular-mass hyaluronan mediates the cancer resistance of the naked mole rat. Nature 499, 346–U122 (2013). 10.1038/nature12234

19 Urriola-Muñoz, P., Pattison, L. A. & Smith, E. S. J. Dysregulation of ADAM10 shedding activity in naked mole-rat fibroblasts is due to deficient phosphatidylserine externalization. Journal of cellular physiology 238, 761–775 (2023). 10.1002/jcp.30972

20 Miyawaki, S. et al. Tumour resistance in induced pluripotent stem cells derived from naked mole-rats. Nature communications 7 (2016). ARTN 11471 10.1038/ncomms11471

21 Oka, K. et al. Resistance to chemical carcinogenesis induction via a dampened inflammatory response in naked mole-rats. Commun Biol 5, 287 (2022). 10.1038/s42003-022-03241-y

22 Hilton, H. G. et al. Single-cell transcriptomics of the naked mole-rat reveals unexpected features of mammalian immunity. PLoS Biol 17, e3000528 (2019). 10.1371/journal.pbio.3000528

23 Lin, T. D. et al. Evolution of T cells in the cancer-resistant naked mole-rat. Nature communications 15 (2024). ARTN 3145 10.1038/s41467-024-47264-x

24 Jamieson, C. H. M. & Weissman, I. L. Stem-Cell Aging and Pathways to Precancer Evolution. New Engl J Med 389, 1310–1319 (2023). 10.1056/NEJMra2304431

25 Stead, E. R. & Bjedov, I. Balancing DNA repair to prevent ageing and cancer. Exp Cell Res 405, 112679 (2021). 10.1016/j.yexcr.2021.112679

26 Page, M. E., Lombard, P., Ng, F., Gottgens, B. & Jensen, K. B. The epidermis comprises autonomous compartments maintained by distinct stem cell populations. Cell stem cell 13, 471–482 (2013). 10.1016/j.stem.2013.07.010

27 Blanpain, C. & Fuchs, E. Epidermal stem cells of the skin. Annu Rev Cell Dev Biol 22, 339–373 (2006). 10.1146/annurev.cellbio.22.010305.104357

28 Matthews, H. K. The cell-division cycle is faster in cell types prone to forming cancer. Nature 641, 1108–1109 (2025). 10.1038/d41586-025-01138-4

29 Braun, K. M. et al. Manipulation of stem cell proliferation and lineage commitment: visualisation of label-retaining cells in wholemounts of mouse epidermis. Development 130, 5241–5255 (2003).

30 Wong, P. & Coulombe, P. A. Loss of keratin 6 (K6) proteins reveals a function for intermediate filaments during wound repair. J Cell Biol 163, 327–337 (2003). 10.1083/jcb.200305032

31 Cockburn, K. et al. Gradual differentiation uncoupled from cell cycle exit generates heterogeneity in the epidermal stem cell layer. Nat Cell Biol (2022). 10.1038/s41556-022-01021-8

32 Cohen, E., Johnson, C., Redmond, C. J., Nair, R. R. & Coulombe, P. A. Revisiting the significance of keratin expression in complex epithelia. J Cell Sci 135 (2022). 10.1242/jcs.260594

33 Aragona, M. et al. Mechanisms of stretch-mediated skin expansion at single-cell resolution. Nature 584, 268-+ (2020). 10.1038/s41586-020-2555-7

34 Wang, S. et al. Single cell transcriptomics of human epidermis identifies basal stem cell transition states. Nat Commun 11, 4239 (2020). 10.1038/s41467-020-18075-7

35 Lin, Z. G. et al. Murine interfollicular epidermal differentiation is gradualistic with GRHL3 controlling progression from stem to transition cell states. Nature communications 11 (2020). ARTN 5434 10.1038/s41467-020-19234-6

36 Moore, R. J. et al. Mice deficient in tumor necrosis factor-alpha are resistant to skin carcinogenesis. Nat Med 5, 828–831 (1999). 10.1038/10552

37 Lin, Y. et al. Role of mammary epithelial and stromal P450 enzymes in the clearance and metabolic activation of 7,12-dimethylbenz(a)anthracene in mice. Toxicol Lett 212, 97–105 (2012). 10.1016/j.toxlet.2012.05.005

38 Ishizuka, T. et al. Human carboxymethylenebutenolidase as a bioactivating hydrolase of olmesartan medoxomil in liver and intestine. J Biol Chem 285, 11892–11902 (2010). 10.1074/jbc.M109.072629

39 Scully, R. et al. Dynamic changes of BRCA1 subnuclear location and phosphorylation state are initiated by DNA damage. Cell 90, 425–435 (1997). 10.1016/s0092-8674(00)80503-6

40 Malak, M. et al. Insights into metabolic changes during epidermal differentiation as revealed by multiphoton microscopy with fluorescence lifetime imaging. Scientific reports 15, 6377 (2025). 10.1038/s41598-025-90101-4

41 Kobayashi, T., Naik, S. & Nagao, K. Choreographing Immunity in the Skin Epithelial Barrier. Immunity 50, 552–565 (2019). 10.1016/j.immuni.2019.02.023

42 Duong, K. L. et al. Identification of hematopoietic-specific regulatory elements from the CD45 gene and use for lentiviral tracking of transplanted cells. Exp Hematol 42, 761–772 e761-710 (2014). 10.1016/j.exphem.2014.05.005

43 Sun, L., Su, Y., Jiao, A., Wang, X. & Zhang, B. T cells in health and disease. Signal Transduct Target Ther 8, 235 (2023). 10.1038/s41392-023-01471-y

44 Sanchez Sanchez, G., et al. Invariant gammadeltaTCR natural killer-like effector T cells in the naked mole-rat. Nat Commun 15, 4248 (2024). 10.1038/s41467-024-48652-z

45 Ma, W. & Zhou, S. Metabolic Rewiring in the Face of Genomic Assault: Integrating DNA Damage Response and Cellular Metabolism. Biomolecules 15 (2025). 10.3390/biom15020168

46 Abel, E. L., Angel, J. M., Kiguchi, K. & DiGiovanni, J. Multi-stage chemical carcinogenesis in mouse skin: fundamentals and applications. Nat Protoc 4, 1350–1362 (2009). 10.1038/nprot.2009.120

47 Krieg, L., Kuhlmann, I. & Marks, F. Effect of tumor-promoting phorbol esters and of acetic acid on mechanisms controlling DNA synthesis and mitosis (Chalones) and on the biosynthesis of histidine-rich protein in mouse epidermis. Cancer Res 34, 3135–3146 (1974).

48 Ellis, P. et al. Reliable detection of somatic mutations in solid tissues by laser-capture microdissection and low-input DNA sequencing. Nat Protoc 16, 841–871 (2021). 10.1038/s41596-020-00437-6

49 de Visser, K. E. & Joyce, J. A. The evolving tumor microenvironment: From cancer initiation to metastatic outgrowth. Cancer cell 41, 374–403 (2023). 10.1016/j.ccell.2023.02.016

50 Cheng, P. S. W., Zaccaria, M. & Biffi, G. Functional heterogeneity of fibroblasts in primary tumors and metastases. Trends Cancer 11, 135–153 (2025). 10.1016/j.trecan.2024.11.005

51 Faubert, B., Solmonson, A. & DeBerardinis, R. J. Metabolic reprogramming and cancer progression. Science 368 (2020). 10.1126/science.aaw5473

52 Zheng, X. et al. ANGPTL4-Mediated Promotion of Glycolysis Facilitates the Colonization of Fusobacterium nucleatum in Colorectal Cancer. Cancer Res 81, 6157–6170 (2021). 10.1158/0008-5472.CAN-21-2273

53 Zhu, C. et al. Adipose-derived stem cells promote glycolysis and peritoneal metastasis via TGF-beta1/SMAD3/ANGPTL4 axis in colorectal cancer. Cell Mol Life Sci 81, 189 (2024). 10.1007/s00018-024-05215-1

54 Ma, Z. et al. CXCL12 alone is enough to Reprogram Normal Fibroblasts into Cancer-Associated Fibroblasts. Cell Death Discov 11, 156 (2025). 10.1038/s41420-025-02420-0

55 Fu, Y. et al. The reverse Warburg effect is likely to be an Achilles’ heel of cancer that can be exploited for cancer therapy. Oncotarget 8, 57813–57825 (2017). 10.18632/oncotarget.18175

56 Martincorena, I. et al. Tumor evolution. High burden and pervasive positive selection of somatic mutations in normal human skin. Science 348, 880–886 (2015). 10.1126/science.aaa6806

57 Naxerova, K. Clonal competition in a confined space. Nat Genet 52, 553–554 (2020). 10.1038/s41588-020-0638-x

58 Clayton, E. et al. A single type of progenitor cell maintains normal epidermis. Nature 446, 185–189 (2007).

59 Alcolea, M. P. et al. Differentiation imbalance in single oesophageal progenitor cells causes clonal immortalization and field change. Nat Cell Biol 16, 615–622 (2014). 10.1038/ncb2963

60 Zhang, S. et al. Tumor initiation and early tumorigenesis: molecular mechanisms and interventional targets. Signal Transduct Target Ther 9, 149 (2024). 10.1038/s41392-024-01848-7

61 Kato, T. et al. Dynamic stem cell selection safeguards the genomic integrity of the epidermis. Dev Cell 56, 3309–3320 e3305 (2021). 10.1016/j.devcel.2021.11.018

62 Stewart, K. S. et al. Stem cells tightly regulate dead cell clearance to maintain tissue fitness. Nature 633, 407–416 (2024). 10.1038/s41586-024-07855-6

63 Park, S. et al. Skin-resident immune cells actively coordinate their distribution with epidermal cells during homeostasis. Nat Cell Biol 23, 476–484 (2021). 10.1038/s41556-021-00670-5

64 Sousa, C. M. et al. Pancreatic stellate cells support tumour metabolism through autophagic alanine secretion. Nature 536, 479–483 (2016). 10.1038/nature19084

65 Huang, K. et al. Tumor metabolic regulators: key drivers of metabolic reprogramming and the promising targets in cancer therapy. Mol Cancer 24, 7 (2025). 10.1186/s12943-024-02205-6

66 Pascual, G. et al. Targeting metastasis-initiating cells through the fatty acid receptor CD36. Nature 541, 41–45 (2017). 10.1038/nature20791

67 Hafemeister, C. & Satija, R. Normalization and variance stabilization of single-cell RNA-seq data using regularized negative binomial regression. Genome Biol 20, 296 (2019). 10.1186/s13059-019-1874-1

68 Satija, R., Farrell, J. A., Gennert, D., Schier, A. F. & Regev, A. Spatial reconstruction of single-cell gene expression data. Nat Biotechnol 33, 495–502 (2015). 10.1038/nbt.3192

69 Lun, A. T., McCarthy, D. J. & Marioni, J. C. A step-by-step workflow for low-level analysis of single-cell RNA-seq data with Bioconductor. F1000Res 5, 2122 (2016). 10.12688/f1000research.9501.2

70 Tarashansky, A. J., Xue, Y., Li, P., Quake, S. R. & Wang, B. Self-assembling manifolds in single-cell RNA sequencing data. eLife 8 (2019). 10.7554/eLife.48994

71 Tarashansky, A. J. et al. Mapping single-cell atlases throughout Metazoa unravels cell type evolution. eLife 10 (2021). 10.7554/eLife.66747

72 Joost, S. et al. The Molecular Anatomy of Mouse Skin during Hair Growth and Rest. Cell stem cell 26, 441–457 e447 (2020). 10.1016/j.stem.2020.01.012

73 Anders, S. & Huber, W. Differential expression analysis for sequence count data. Genome Biol 11, R106 (2010). gb-2010-11-10-r106 [pii] 10.1186/gb-2010-11-10-r106

74 Subramanian, A. et al. Gene set enrichment analysis: a knowledge-based approach for interpreting genome-wide expression profiles. Proc Natl Acad Sci U S A 102, 15545–15550 (2005). 10.1073/pnas.0506580102

75 Liberzon, A. et al. The Molecular Signatures Database (MSigDB) hallmark gene set collection. Cell Syst 1, 417–425 (2015). 10.1016/j.cels.2015.12.004

76 Milacic, M. et al. The Reactome Pathway Knowledgebase 2024. Nucleic Acids Res 52, D672–D678 (2024). 10.1093/nar/gkad1025

77 Kanehisa, M., Furumichi, M., Sato, Y., Matsuura, Y. & Ishiguro-Watanabe, M. KEGG: biological systems database as a model of the real world. Nucleic Acids Res 53, D672–D677 (2025). 10.1093/nar/gkae909

78 Aibar, S. et al. Identification of expression patterns in the progression of disease stages by integration of transcriptomic data. BMC Bioinformatics 17, 432 (2016). 10.1186/s12859-016-1290-4

79 Kawamura, Y. et al. Cellular senescence induction leads to progressive cell death via the INK4a-RB pathway in naked mole-rats. Embo J 42, e111133 (2023). 10.15252/embj.2022111133

80 Schubert, M. et al. Perturbation-response genes reveal signaling footprints in cancer gene expression. Nat Commun 9, 20 (2018). 10.1038/s41467-017-02391-6

81 Rognoni, E. et al. Role of distinct fibroblast lineages and immune cells in dermal repair following UV radiation-induced tissue damage. eLife 10 (2021). 10.7554/eLife.71052

82 Badia, I. M. P. et al. decoupleR: ensemble of computational methods to infer biological activities from omics data. Bioinform Adv 2, vbac016 (2022). 10.1093/bioadv/vbac016

83 Armingol, E., Officer, A., Harismendy, O. & Lewis, N. E. Deciphering cell-cell interactions and communication from gene expression. Nat Rev Genet 22, 71–88 (2021). 10.1038/s41576-020-00292-x

84 Dimitrov, D. et al. Comparison of methods and resources for cell-cell communication inference from single-cell RNA-Seq data. Nat Commun 13, 3224 (2022). 10.1038/s41467-022-30755-0

85 Bates, D., Mächler, M., Bolker, B. & Walker, S. Fitting Linear Mixed-Effects Models Using lme4. (2015). 10.18637/jss.v067.i01

86 Li, H. & Durbin, R. Fast and accurate short read alignment with Burrows-Wheeler transform. Bioinformatics 25, 1754–1760 (2009). 10.1093/bioinformatics/btp324

87 Danecek, P. et al. Twelve years of SAMtools and BCFtools. Gigascience 10 (2021). 10.1093/gigascience/giab008

88 McKenna, A. et al. The Genome Analysis Toolkit: a MapReduce framework for analyzing next-generation DNA sequencing data. Genome Res 20, 1297–1303 (2010). 10.1101/gr.107524.110

89 McLaren, W. et al. The Ensembl Variant Effect Predictor. Genome Biol 17, 122 (2016). 10.1186/s13059-016-0974-4

90 Sondka, Z. et al. The COSMIC Cancer Gene Census: describing genetic dysfunction across all human cancers. Nat Rev Cancer 18, 696–705 (2018). 10.1038/s41568-018-0060-1

